# RNA-seq Analysis Identifies Host Pathways Linked to Polyphosphate-Mediated Macrophage Dysfunction

**DOI:** 10.64898/2026.11.25.769283

**Authors:** Anushka Tennakoon, Madelyn Moresi, Richard H. Gomer, Ramesh Rijal

## Abstract

After ingestion into macrophage phagosomes, *Mycobacterium tuberculosis* (Mtb) evades killing by preventing phagosome acidification and fusion of the phagosome with a lysosome. Mtb accumulates extracellular polyphosphate (polyP), and polyP inhibits macrophage phagosome acidification, lysosome activity, and bacterial killing. Whether polyP alters macrophage gene expression to accomplish this remains unclear. Here, we performed RNA sequencing on primary human monocyte-derived macrophages treated with exogenous polyP or infected with attenuated Mtb. Overall, macrophage gene expression clustered primarily by donor sex and individual identity. Mtb altered levels of 513 mRNAs in female macrophages compared to 35 mRNAs in male macrophages, predominantly downregulating transcripts in females, whereas polyP altered a smaller subset of mRNAs in both sexes. Both polyP and Mtb altered gene ontology pathways associated with cytoplasmic translation, ribosomal subunits, and adaptive immune responses. Quantitative reverse transcription-PCR validation of a candidate-gene panel showed that Mtb downregulated *CORO1A* in both sexes and polyP downregulated *CORO1A* in females, whereas both polyP and Mtb downregulated *S1PR1* and *INPP4B* in male and female macrophages. Mtb also increased *CLN8* mRNA in female macrophages without altering total CLN8 protein as assessed by immunofluorescence. Mtb alters host lipid metabolism to induce the formation of lipid-laden foamy macrophages, which serve as a nutrient-rich reservoir that supports intracellular Mtb persistence. PolyP increased neutral lipid accumulation in male and female macrophages, whereas Mtb increased neutral lipid accumulation in female macrophages. Together, these findings identify macrophage transcriptional and lipid pathways modulated by polyP and Mtb.

**Importance:** Human macrophages engulf bacteria into phagosomes, which then fuse with lysosomes to kill the bacteria. However, after engulfment, *Mycobacterium tuberculosis* (Mtb) can block phagosome-lysosome fusion, allowing its intracellular survival. We previously found that Mtb secretes extracellular polyphosphate (polyP) to inhibit phagosome maturation and bacterial killing. Here, we show that Mtb and exogenous polyP alter macrophage expression of genes involved in translation, adaptive immunity, endolysosomal trafficking, and promote neutral lipid accumulation. These findings identify host pathways modulated by bacterial polyP that may inform host-directed strategies against tuberculosis.

## Introduction

Polyphosphate (polyP) is a linear chain of phosphate units found across all forms of life, and polyP metabolism is closely associated with the virulence of *Mycobacterium tuberculosis* (Mtb) and other intracellular pathogens (1–4). Macrophages normally engulf invading microorganisms into a membrane-bound compartment called the phagosome, which then acidifies and fuses with lysosomes to form a phagolysosome, wherein hydrolytic enzymes and antimicrobial compounds kill the ingested pathogen (5–7). However, Mtb causes an estimated 1.23 million global tuberculosis deaths annually (8), in part because Mtb can prevent the fusion of phagosomes with lysosomes, evade bactericidal actions, and persist within macrophages (9, 10). Mtb accumulates extracellular polyP, and this polyP inhibits macrophage phagosome acidification, lysosome activity, and bacterial killing (11). Conversely, treatment of Mtb-infected macrophages with the polyP-degrading enzyme exopolyphosphatase (PPX) reduces the survival of phagocytosed Mtb (11).

The mechanisms by which polyP can inhibit macrophage phagosome maturation involve evolutionarily conserved signal transduction pathways (12). In the eukaryotic microbe and professional phagocyte *Dictyostelium discoideum*, polyP requires the G protein-coupled receptor glutamate receptor-like protein D (GrlD), the GPCR-interacting arrestin-like protein AdcB, inositol hexakisphosphate kinase A (I6kA), the Rho GTPase RacE, and the target of rapamycin (TOR) component Lst8 to inhibit the killing of ingested bacteria (12). Pharmacological inhibition of human orthologs of these proteins, including mammalian target of rapamycin (mTOR) and inositol hexakisphosphate kinase (IP6K), as well as polyP-sensing receptors such as the purinergic receptor P2Y1 and the receptor for advanced glycation end products (RAGE), potentiates the ability of macrophages to kill Mtb (13). Although transcriptomic studies have identified host genes that are differentially expressed during broad Mtb infection (14), the downstream transcriptional consequences of polyP-induced signaling in macrophages remain uncharacterized. Because TB incidence is higher in males than in females, (15, 16), sex-specific macrophage biology must be accounted for when evaluating these host-pathogen interactions. Therefore, comparing the transcriptional signatures elicited by polyP and Mtb, while accounting for established sex biases, is necessary to identify the specific host genes targeted by polyP to arrest phagosome maturation.

In this report, we performed RNA-sequencing on primary human monocyte-derived macrophages from male and female donors to compare the transcriptional signatures elicited by exogenous bacterial polyP and Mtb infection. We found that polyP and Mtb induce macrophage transcriptomic responses that cluster primarily by sex, with Mtb inducing a substantially larger transcriptional response in female donors. Using a candidate gene panel, we identified and validated altered expression of genes involved in phagosome maturation and lysosomal function. PolyP and Mtb also increased neutral-lipid accumulation in human macrophages. Together, these findings suggest a set of candidate genes involved in macrophage responses to polyP. Future studies will be needed to determine how these transcriptional changes vary between sexes and whether they affect the clearance of intracellular pathogens.

## Results

### PolyP and Mtb induce macrophage transcriptomic responses that cluster primarily by sex

Mycobacteria accumulate extracellular polyP that inhibit phagosome acidification, lysosome activity, and potentiates the survival of ingested bacteria in human macrophages (11). Pharmacological inhibitors of host proteins that may mediate polyP responses potentiate the ability of macrophages to kill ingested Mtb, *Legionella*, and *Listeria* (13). However, how polyP alters macrophage gene expression, and how this compares with the response to Mtb is unclear. To address this, human blood monocytes from three male and three female donors were cultured with granulocyte-macrophage colony-stimulating factor (GM-CSF) for 6 days to generate macrophages (17). The macrophages were then left untreated (control), treated with 15 µg/mL polyP, or infected with Mtb, at 5 approximately bacteria per macrophage, for 24 hours, and RNA was analyzed by RNA sequencing (RNA-seq), with selected genes validated by qRT-PCR.

In a principal component analysis (PCA) (18), principal component 1 (PC1; 44% of the variance) separated samples from male and female donors, and PC2 (29% of the variance) separated samples from one male donor from those of the other donors (**Figure 1A**). Samples from each donor clustered together regardless of treatment, indicating that differences between sexes and between donors were larger than the effects of polyP or Mtb. Within each donor cluster, however, Mtb-infected samples were consistently displaced from control samples, whereas polyP-treated samples were generally positioned closer to the controls. Hierarchical clustering of sample-to-sample Euclidean distances also broadly separated samples by sex, consistent with the PCA (**Figure S1A**). Samples had 9.1 ± 1.1 million mapped reads (mean ± SD, n = 18), all samples yielded more than 5 million mapped reads, with similar sequencing depths across treatments and sexes (**Figure 1B**; **Figure S1B**). In each sample, 43.2 ± 1.4% of annotated genes were detected (**Figure 1C**), and the 100 most highly expressed genes accounted for 34.5 ± 1.6% of mapped reads (**Figure S1C**), indicating comparable library complexity across samples. Together, these data suggest that the RNA-seq library quality was uniform across samples, and overall gene expression was determined primarily by donor sex and individual identity, which outweighed polyP or Mtb treatment.

**Figure 1.**
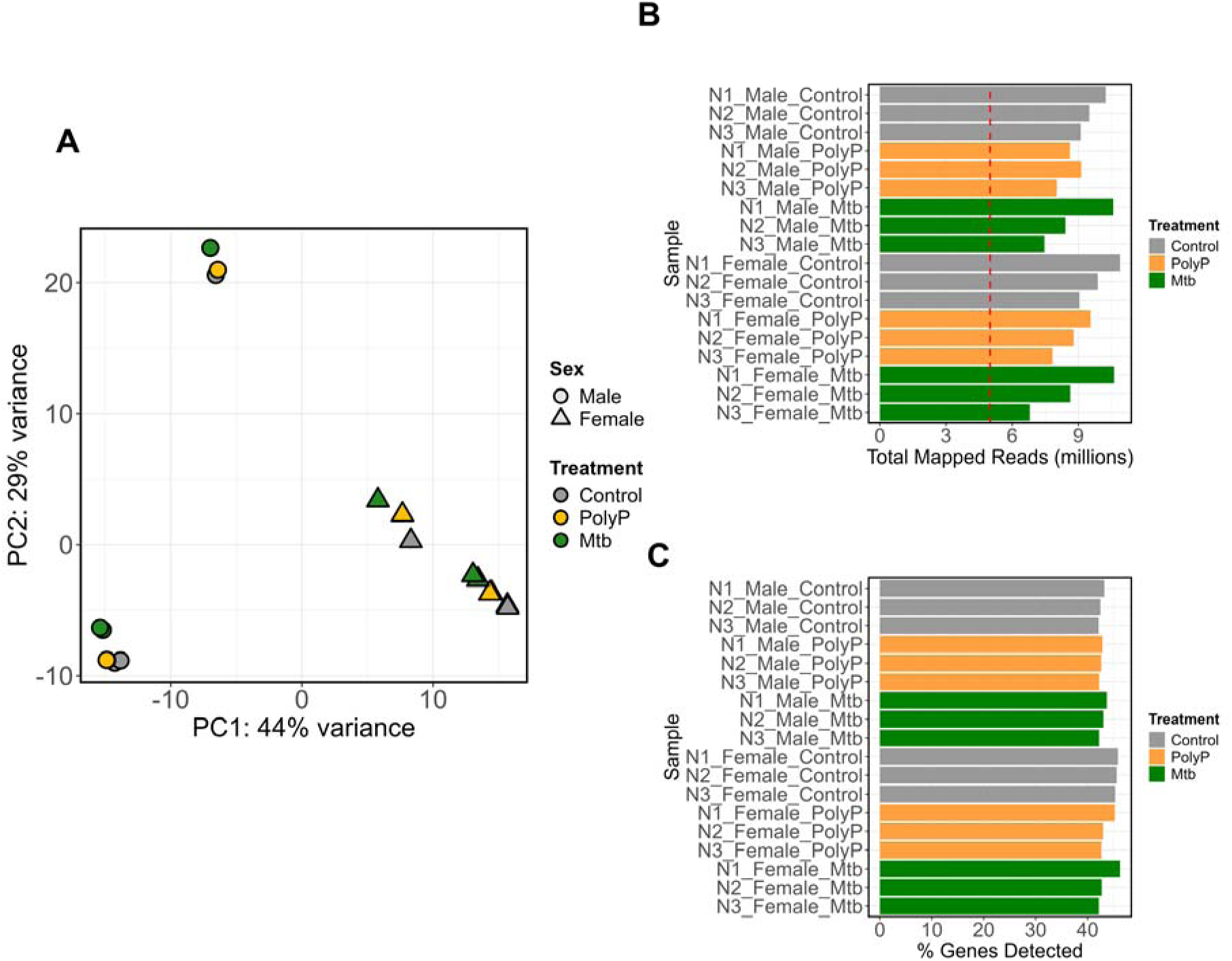
Principal component analysis and sequencing quality metrics. (**A**) Principal component analysis (PCA) plot of gene expression profiles across the 18 macrophages RNA-seq samples from male (circles) and female (triangles) donors under untreated (Control, gray), polyP-treated (PolyP, orange), and Mtb-infected (Mtb, green) conditions, plotted along principal component 1 (PC1; 44% of variance) and PC2 (29% of variance). (**B**) Total mapped reads (in millions) per sample across control (gray), polyP (orange), and Mtb (green) conditions. The red dashed line indicates 5 million mapped reads. (**C**) Percentage of annotated human genes detected per sample across control (gray), polyP (orange), and Mtb (green) conditions.

### PolyP induces a smaller transcriptional response than Mtb

To determine whether polyP and Mtb regulate overlapping sets of macrophage genes, we first compared polyP-treated and Mtb-infected macrophages with untreated/uninfected control in a combined, sex-averaged analysis. mRNAs with an absolute log_2_ fold change >1 and a Benjamini–Hochberg-adjusted P <0.05 were considered differentially expressed. Compared with controls, polyP upregulated 3 mRNAs and downregulated 7 (**Figure 2A**), whereas Mtb upregulated 4 mRNAs and downregulated 323 (**Figure 2B**). *SCARF1* (scavenger receptor class F member 1), encoding a receptor that mediates engulfment of apoptotic cells (19), and *BTBD7* (BTB domain-containing 7), encoding a regulator of E-cadherin turnover and cell motility (20), were upregulated by both treatments, with one mRNA upregulated only by polyP and two only by Mtb (**Figure 2C**). *HSPA1A* (heat shock protein family A member 1A), a stress-inducible Hsp70 chaperone (21); *SELL* (L-selectin), a leukocyte adhesion molecule (22); *S1PR1* (sphingosine-1-phosphate receptor 1), a regulator of lymphocyte egress (23); *KLRD1* (killer cell lectin-like receptor D1, CD94), a component of the NK-cell receptor for HLA-E (24, 25); and *LEF1* (lymphoid enhancer-binding factor 1), a Wnt/β-catenin transcription factor (26), were downregulated by both treatments, with two mRNAs downregulated only by polyP and 318 only by Mtb (**Figure 2D**). The analysis did not find any mRNAs that were upregulated in polyP treatment and downregulated in Mtb treatment or vice versa. Together, these data suggest that polyP causes a small transcriptional response in macrophages that partly overlaps with the much larger response to Mtb.

**Figure 2.**
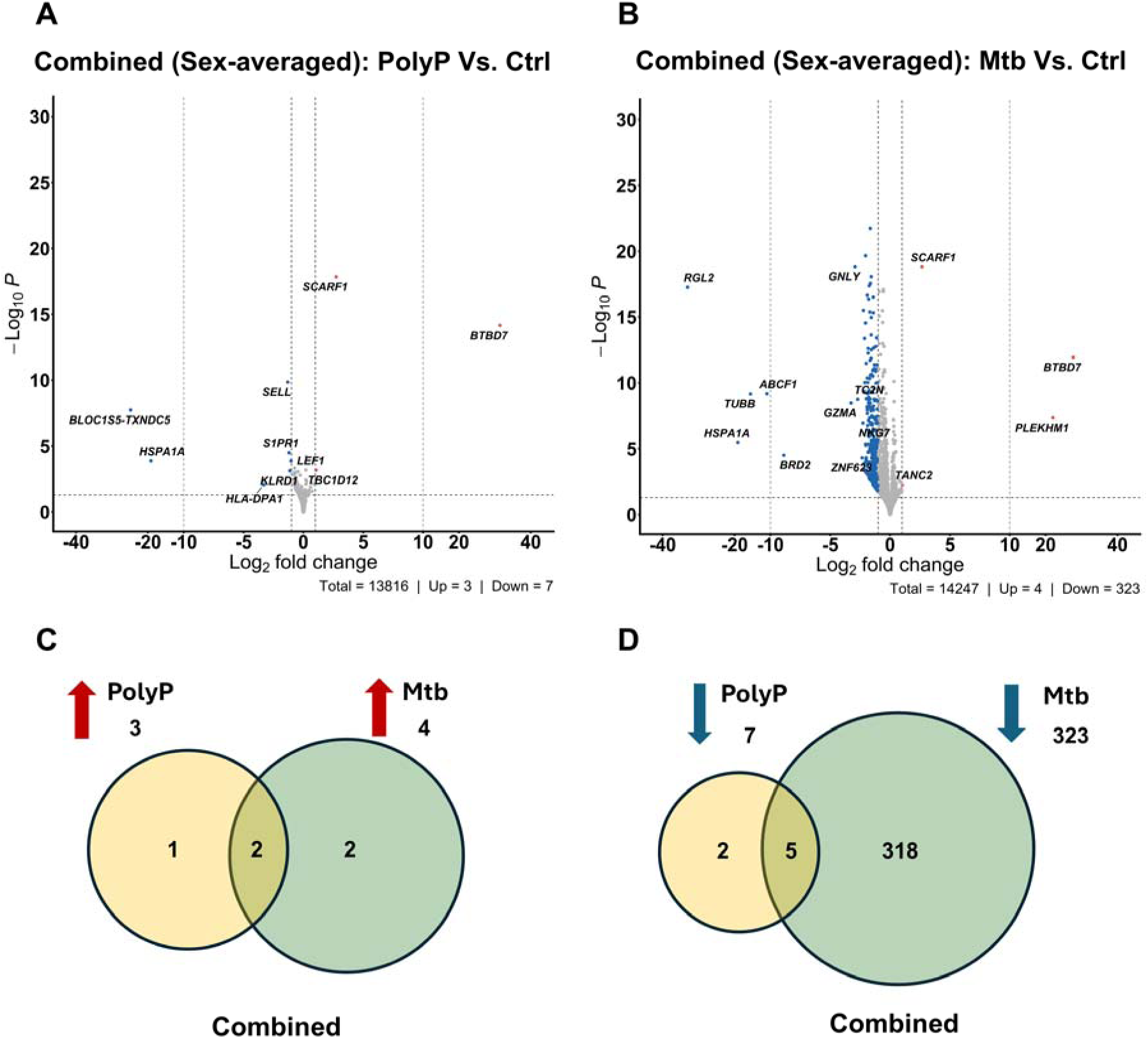
Differential gene expression in the combined sex-averaged RNA-seq analysis. A and. **B**) Volcano plots of differentially expressed genes (DEGs) in the combined (sex-averaged) analysis comparing polyP-treated (**A**) or Mtb-infected (**B**) macrophages with untreated controls (Ctrl). Vertical dashed lines indicate log_2_ fold change = 1, and the horizontal dashed line indicates a Benjamini–Hochberg-adjusted *P* value of 0.05. Upregulated genes are shown in red, downregulated genes are shown in blue, and genes not meeting differential-expression criteria are shown in gray. (**C and D**) Venn diagrams show the number of unique and overlapping genes significantly upregulated (**C**) and downregulated (**D**) in polyP-treated (yellow) and Mtb-infected (green) macrophages compared with controls in the combined analysis.

### PolyP and Mtb both alter levels of macrophage mRNAs associated with translation and adaptive immunity

To compare differentially expressed genes (DEGs) across individual samples, we generated heatmaps of row-scaled (Z-score) expression values, with genes grouped by hierarchical clustering (27). In the polyP comparison, *LEF1*, *SELL*, and *S1PR1* showed the highest expression in female control samples and lower expression in female polyP-treated samples (**Figure 3A**). *HSPA1A* and *BLOC1S5-TXNDC5* (a read-through transcript spanning the neighboring *BLOC1S5* and *TXNDC5* genes) showed the highest expression in one male control sample (**Figure 3A**). Among the genes upregulated by polyP, *SCARF1* showed the lowest expression in male control samples, and *BTBD7* and *TBC1D12* (TBC1 domain family member 12) showed the highest expression in polyP-treated samples (**Figure 3A**). In the Mtb comparison, most DEGs showed the highest expression in female control samples and lower expression in female Mtb-infected samples (**Figure 3B**).

**Figure 3.**
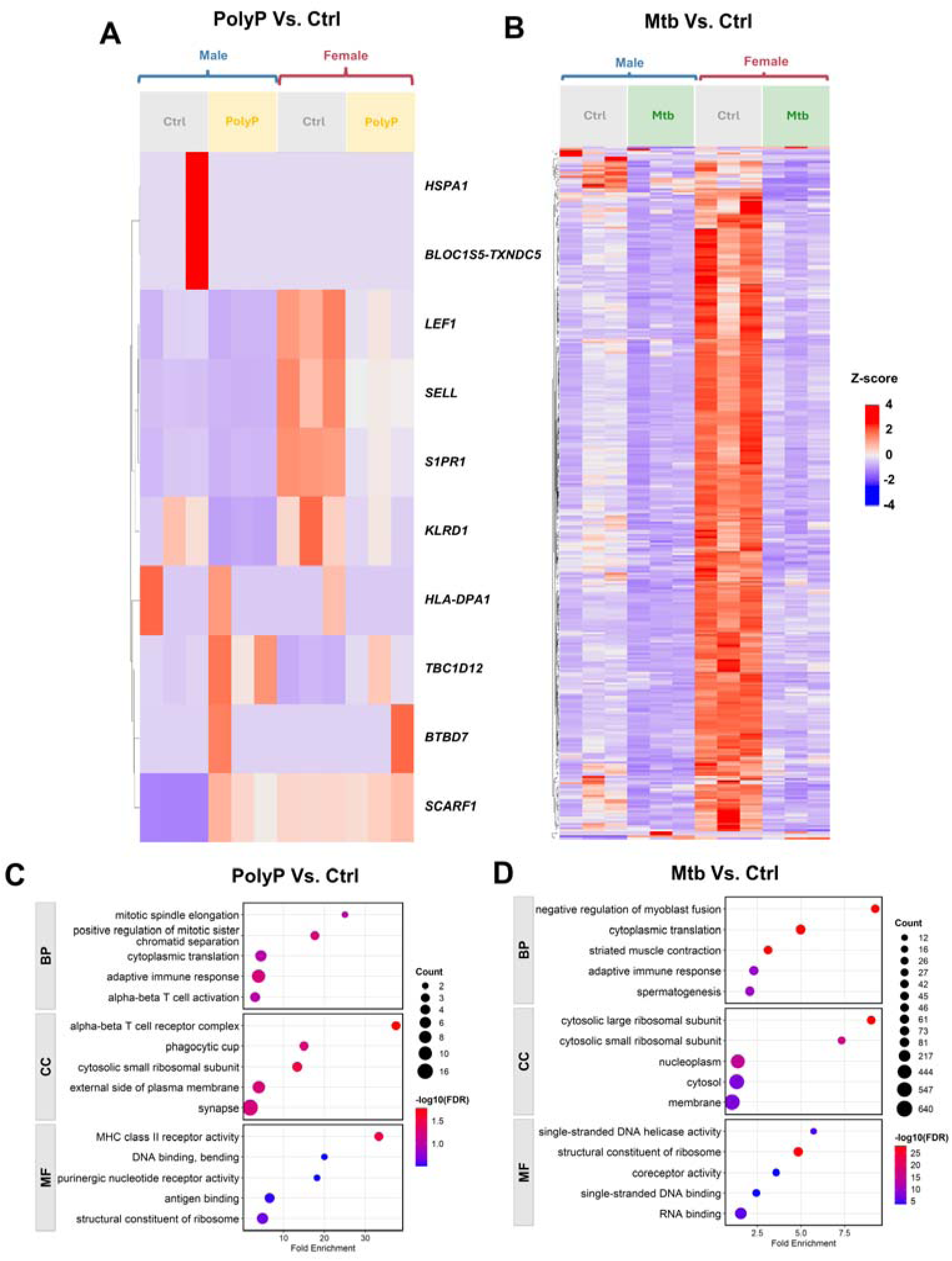
Clustered expression heatmaps and Gene Ontology enrichment. (**A and B**) Heatmaps of row-scaled (*Z*-score) expression values for differentially expressed genes in polyP-treated vs. control (Ctrl) macrophages (A) and Mtb-infected vs. control macrophages (B) across individual male and female donors. (**C and D**) Gene Ontology (GO) enrichment dot plots for biological processes (BP), cellular components (CC), and molecular functions (MF) of genes significantly altered by polyP (**C**) or Mtb (**D**) compared with controls for the combined male and female data. The top five enriched terms in each category are plotted by fold enrichment; dot size indicates the number of enriched genes (Count), and dot color indicates statistical significance expressed as –log10(FDR).

To identify the biological functions associated with the transcriptional changes, we performed GO enrichment analysis using all genes significantly altered by each treatment (adjusted P <0.05). In polyP-treated macrophages, the most enriched biological processes were mitotic spindle elongation, positive regulation of mitotic sister chromatid separation, cytoplasmic translation, adaptive immune response, and alpha-beta T cell activation (**Figure 3C**). The most enriched cellular components were the alpha-beta T cell receptor complex, phagocytic cup, cytosolic small ribosomal subunit, external side of plasma membrane, and synapse. The most enriched molecular functions were MHC class II receptor activity, DNA binding, bending, purinergic nucleotide receptor activity, antigen binding, and structural constituent of ribosome (**Figure 3C**). In Mtb-infected macrophages, the most enriched biological processes were negative regulation of myoblast fusion, cytoplasmic translation, striated muscle contraction, adaptive immune response, and spermatogenesis (**Figure 3D**). The most enriched cellular components were the cytosolic large ribosomal subunit, cytosolic small ribosomal subunit, nucleoplasm, cytosol, and membrane. The most enriched molecular functions were single-stranded DNA helicase activity, structural constituent of ribosome, coreceptor activity, single-stranded DNA binding, and RNA binding (**Figure 3D**). Cytoplasmic translation, adaptive immune response, cytosolic small ribosomal subunit, and structural constituent of ribosome were enriched with both treatments (**Figures 3C and D**). Together, these data suggest that polyP and Mtb both alter the expression of macrophage genes associated with translation and adaptive immunity.

### Sex-stratified analysis reveals a substantially larger transcriptional response to Mtb in female donors

Mtb infection occurrence is higher in males compared to females (16), and studies using mouse models have shown that male mice are more susceptible to Mtb infection than female mice (15). To determine whether macrophage responses to polyP or Mtb differed by donor sex, we performed DEG analysis separately for female and male donors. In response to polyP, one gene was upregulated and five genes were downregulated in female donors, whereas three genes were upregulated and two genes were downregulated in male donors (**Figures 4A and B**). In response to Mtb, 15 genes were upregulated and 498 genes were downregulated in female donors, whereas eight genes were upregulated and 27 genes were downregulated in male donors (**Figures 4C and D**). Thus, relatively few genes met the differential-expression criteria following polyP treatment in either sex. In the sex-stratified analyses, more genes met these criteria following Mtb infection in female donors (513 total: 15 upregulated and 498 downregulated) than in male donors (35 total: 8 upregulated and 27 downregulated). These counts describe the within-sex analyses and do not by themselves establish a statistically significant sex-by-treatment difference. In female donors, *BTBD7* was upregulated by both polyP and Mtb (**Figure 4E**). Of the five genes downregulated by polyP, three, *S1PR1*, *IKZF3*, and *SELL*, were also downregulated by Mtb (**Figure 4F**). In male donors, *SCARF1* and *BTBD7* were upregulated by both polyP and Mtb (**Figure 4G**), whereas none of the genes downregulated by polyP overlapped with those downregulated by Mtb (**Figure 4H**).

**Figure 4.**
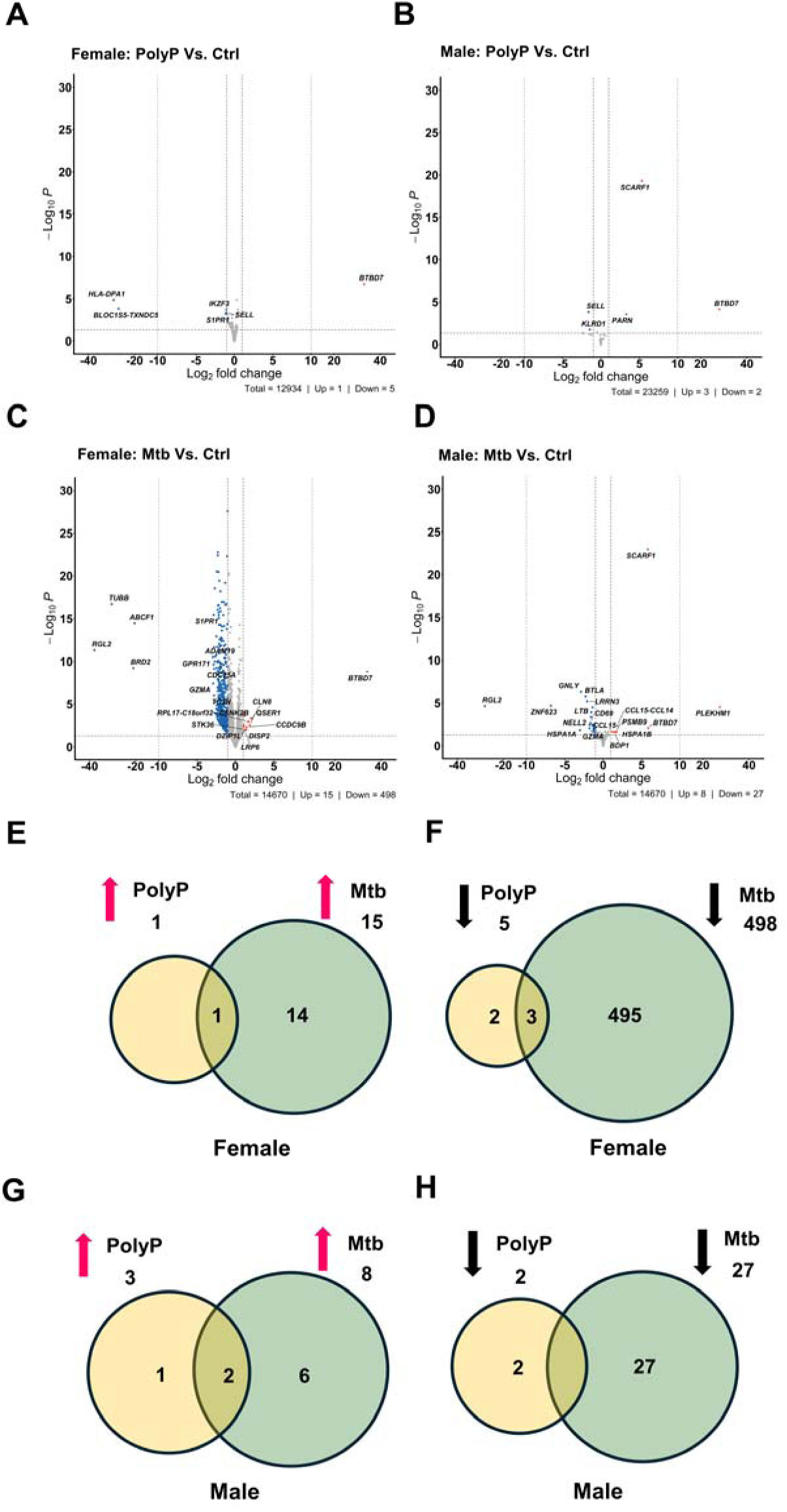
Differential gene expression in the sex-stratified RNA-seq analysis. (**A–D**) Volcano plots of differentially expressed genes comparing polyP-treated vs. control (Ctrl) from female (**A**) and male donors (**B**), or Mtb-infected vs. control macrophages from female (**C**) and male donors (**D**). Vertical dashed lines indicate log_2_ fold change = 1, and horizontal dashed lines indicate a Benjamini–Hochberg-adjusted *P* value of 0.05. Upregulated genes are shown in red, downregulated genes are shown in blue, and non-significant genes are shown in gray. (**E–H**) Venn diagrams showing the number of unique and overlapping genes significantly upregulated (**E and G**) and downregulated (**F and H**) by polyP (yellow) and Mtb (green) compared with controls in female (E and F) and male (G and H) macrophages.

In a heatmap analysis, three of the genes (*S1PR1*, *SELL*, and *IKZF3*) altered by polyP in female donors showed a consistent decreased expression pattern, whereas two genes (*HLA-DPA1* and *BTBD7*) were driven by high expression in a single control sample, and the single upregulated gene (*BTBD7*) was highly expressed in only one polyP-treated sample (**Figures S2A**). Furthermore, most genes differentially expressed following Mtb infection in female donors had higher expression in control samples and lower expression following infection, consistent with the overwhelming predominance of downregulated genes. In contrast, only a subset of genes (15 genes) was upregulated following Mtb infection in females (**Figure S2B**). GO analysis of genes altered by polyP in female donors identified biological processes associated with mitotic regulation and RNA polymerase II-dependent transcription (**Figure S3A**). Genes altered by Mtb in female donors were associated with cytoplasmic translation, mitotic DNA replication, cytosolic ribosomal subunits, and RNA- and single-stranded DNA-binding functions (**Figure S3B**). In male donors, polyP and Mtb altered smaller sets of genes compared to female donors, and several of these changes reflected high expression in individual donors (**Figures S2C and D**). Regulation of immune response biological processes enriched GO terms were detected following polyP treatment in male donors (**Figure S3C**), whereas genes altered by Mtb in male donors were associated with immune response and cytolytic granules (**Figure S3D**). Together, these data suggest that Mtb induces a broad transcriptional response in female macrophages that affects translation and DNA replication, whereas the smaller response in male macrophages affects a limited set of immune-related genes.

### A candidate-gene panel identifies lysosomal, phagosomal, and immune-receptor responses

To further investigate how polyP and Mtb affect macrophage functions, we selected eight genes that showed treatment-associated expression changes and represented pathways implicated by our previous studies, including phagosome maturation, lysosomal function, polyP-responsive signaling, proteasome activity, and antigen presentation. Coronin 1 regulates actin-dependent phagosome maturation (28), and polyP causes coronin 1 retention on macrophage phagosomes (11). *CLN8* (CLN8 transmembrane ER and ERGIC protein) facilitates the transport of lysosomal enzymes and is required for lysosome biogenesis (29), whereas *PLEKHM1* (pleckstrin homology and RUN domain containing M1) promotes late endosome- and autophagosome–lysosome fusion (30). PolyP inhibits phagosome acidification and lysosomal activity (11). *INPP4B* (inositol polyphosphate-4-phosphatase type II B) regulates phosphoinositide-dependent endolysosomal trafficking (31) and was selected together with *S1PR1* because our previous work implicated IP6K- and Akt–mTOR-associated signaling in macrophage responses to polyP (13). *PSMB9* (proteasome 20S subunit beta 9) is an immunoproteasome subunit involved in generating peptides for MHC class I antigen presentation (32) and polyP and Mtb inhibit proteasome activity in macrophages (11). *HLA-DPA1* (major histocompatibility complex, class II, DP alpha 1) was selected to represent MHC class II-mediated antigen presentation (33), another immune function affected during Mtb infection (34). *SCARF1*, which participates in phagocytic clearance of bacteria and apoptotic cells (19), was upregulated by both polyP and Mtb. Together, these eight genes provided a focused panel linking the RNA-seq results to macrophage pathways affected by polyP and Mtb (**Table 1**) (19, 29, 30, 32, 33, 35–37).

**Table 1.** Candidate genes selected from the RNA-seq and RT-PCR analysis, their direction of regulation in male and female macrophages, and their reported functions.

|  |  | **Male** |  | **Female** |  |  |  |
|---|---|---|---|---|---|---|---|
| **Gene Symbol** | **Full Name** | **Regulation in polyP** | **Regulation in Mtb** | **Regulation in polyP** | **Regulation in Mtb** | **Function** | **Reference** |
| ***CORO1A*** | Coronin-1A |  | Downregulated | Downregulated | Downregulated | *CORO1A* inhibits autophagosome formation in alveolar macrophages, thereby facilitating *M. tuberculosis* survival within the lung | (Seto et al., 2012) |
| ***PLEKHM1*** | Pleckstrin homology domain-containing family M member 1 |  | Upregulated |  |  | Rab7/Arl8b-associated adaptor that promotes late endosome and autophagosome–lysosome fusion | (Marwaha et al., 2017) |
| ***INPP4B*** | Inositol polyphosphate-4-phosphatase type II B | Downregulated | Downregulated | Downregulated | Downregulated | Phosphoinositide phosphatase with connection to late endosome formation, endolysosomal trafficking, and lysosome-directed phosphoinositide conversion. | (Rodgers et al., 2021) |
| ***PSMB9*** | Proteasome 20S Subunit Beta 9 |  | Upregulated |  |  | Involved in antigen processing to generate class I binding peptides. | (Kasahara & Flajnik, 2019) |
| ***CLN8*** | CLN8 ER-Lysosome Membrane Linker |  |  |  | Upregulated | ER-resident cargo receptor facilitating transport of lysosomal enzymes, essential for lysosome biogenesis | (di Ronza et al., 2018) |
| ***SCARF1*** | Scavenger Receptor Class F Member 1 | Upregulated | Upregulated |  |  | Mediates bacterial clearance and apoptotic cell removal | (Patten, 2018) |
| ***S1PR1*** | Sphingosine-1-phosphate receptor 1 | Downregulated | Downregulated | Downregulated | Downregulated | Induction of antimicrobial activity in human macrophages that leads to the intracellular killing of nonpathogenic *Mycobacterium smegmatis* and pathogenic *M. tuberculosis* | (Garg et al., 2004) |
| ***HLA-DPA1*** | Major histocompatibility complex, class II, DP alpha 1 |  |  | Downregulated |  | Plays a central role in the immune system by presenting peptides derived from extracellular proteins | (Yamashita et al., 2017) |

We generated a heatmap to visualize the expression patterns and clustering of the eight candidate genes. The genes separated into three clusters: cluster 1 contained *SCARF1* and *CLN8*; cluster 2 contained *HLA-DPA1, INPP4B, CORO1A,* and *S1PR1*; and cluster 3 contained *PLEKHM1* and *PSMB9* (**Figure 5A**). In female donors, *INPP4B, CORO1A,* and *S1PR1* showed higher expression in control samples than in polyP-treated or Mtb-infected samples, with expression generally lowest following Mtb infection. This treatment-associated pattern was less apparent in male donors, although *INPP4B* was downregulated by Mtb in both female and male donors.

**Figure 5.**
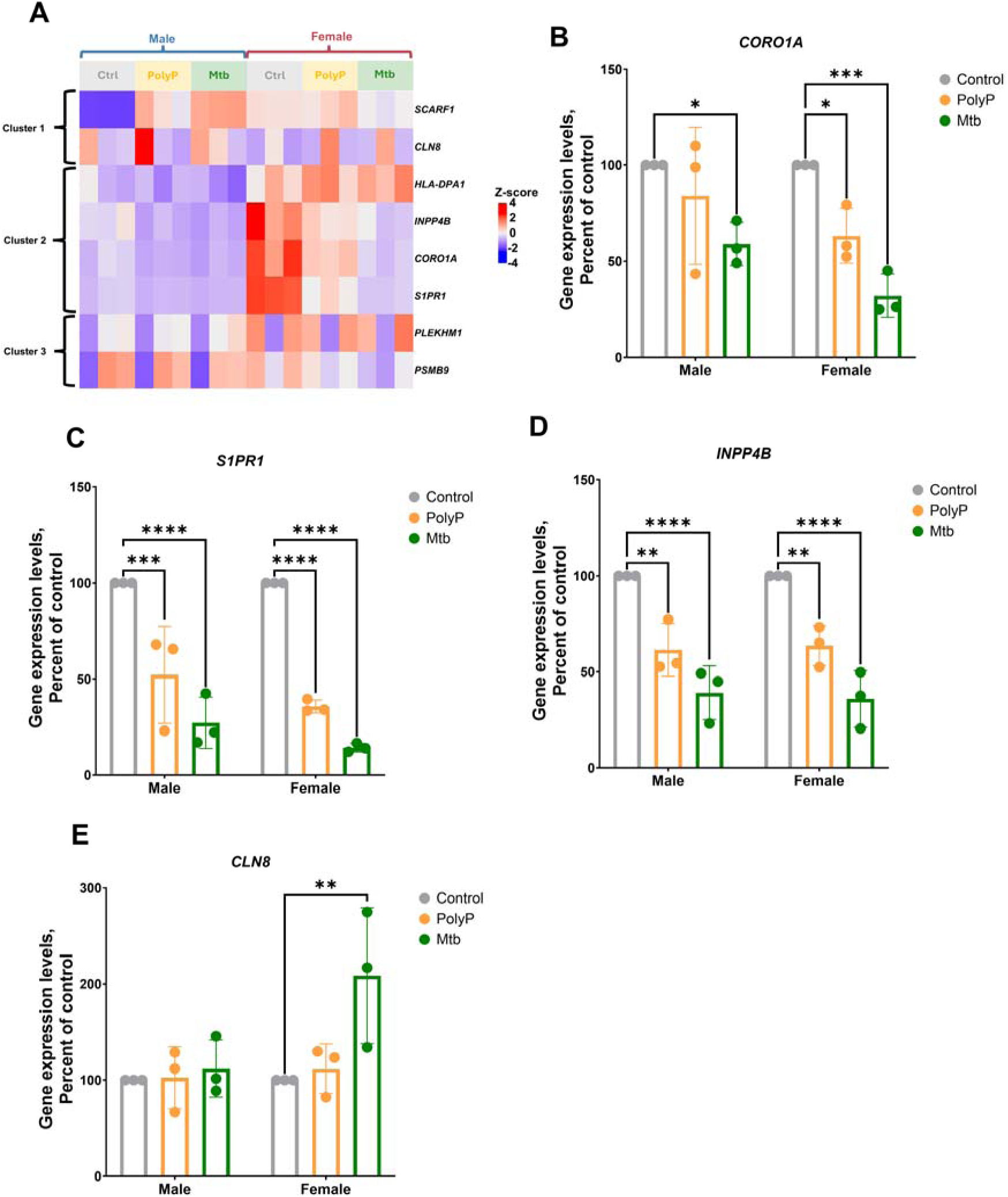
Candidate-gene expression clustering and qRT-PCR validation. (**A**) Heatmap of row-scaled (*Z*-score) RNA-seq expression values for eight selected candidate genes across untreated (Ctrl), polyP-treated (PolyP), and Mtb-infected (Mtb) macrophages from male and female donors, grouped into three hierarchical clusters (Clusters 1–3). (**B–E**) Relative mRNA expression levels of *CORO1A* (**B**), *S1PR1* (**C**), *INPP4B* (**D**), and *CLN8* (**E**) in untreated (Control, gray), polyP-treated (PolyP, orange), and Mtb-infected (Mtb, green) macrophages from male and female donors at 24 hours, determined by qRT-PCR and normalized to *GAPDH*. For each experiment with each donor, expression in the control was set to 100%. All values are mean ± SEM of three male and three female donors. *\*\**p < 0.01; ***p < 0.001; ****p < 0.0001 compared to the control (two-way analysis of variance with Dunnett’s test).

We next used quantitative reverse transcription-PCR (qRT-PCR) to validate the RNA-seq expression patterns of the eight candidate genes. The RNA-seq patterns of four genes, *CORO1A, S1PR1, INPP4B,* and *CLN8*, were reproduced by qRT-PCR. *CORO1A* was downregulated by Mtb in macrophages from both female and male donors. PolyP also downregulated *CORO1A* in macrophages from female donors (**Figure 5B**). *S1PR1* was downregulated by polyP and Mtb in macrophages from both female and male donors, extending the RNA-seq results, in which significant downregulation was detected only in female donors (**Figure 5C**). *INPP4B* was downregulated by Mtb in both female and male donors, consistent with the RNA-seq results, and qRT-PCR additionally showed downregulation by polyP in both sexes (**Figure 5D**). Mtb increased CLN8 expression in macrophages from female donors, consistent with the RNA-seq results (**Figure 5E**). The qRT-PCR expression patterns of *PSMB9, HLA-DPA1, PLEKHM1,* and *SCARF1* did not reproduce the corresponding RNA-seq results (**Figures S4A–D**). Expression of the reference gene *GAPDH* (glyceraldehyde-3-phosphate dehydrogenase) did not differ significantly among the treatment groups (**Figure S4E**).

### PolyP and Mtb promote lipid accumulation in human macrophages

The qRT-PCR analysis identified *CLN8* as a candidate whose expression was increased by Mtb in macrophages from female donors (**Figure 5E**). CLN8 is an endoplasmic reticulum and ER–Golgi intermediate compartment cargo receptor that facilitates the transport of newly synthesized lysosomal enzymes from the endoplasmic reticulum to the Golgi and supports lysosome biogenesis (29, 38). Because lysosomal function and lipid metabolism contribute to macrophage responses during Mtb infection (39, 40), we examined CLN8 protein abundance and neutral-lipid accumulation in macrophages from male and female donors following polyP treatment or Mtb infection.

To determine whether the increase in *CLN8* mRNA was accompanied by a change in CLN8 protein abundance, macrophages were stained with an antibody against CLN8 (**Figure 6A**). CLN8 staining did not differ significantly among untreated, polyP-treated, and Mtb-infected macrophages from either male or female donors (**Figure 6B**). These results indicate that the increase in *CLN8* mRNA detected in female Mtb-infected macrophages was not accompanied by a detectable change in total CLN8 immunofluorescence under these conditions.

**Figure 6.**
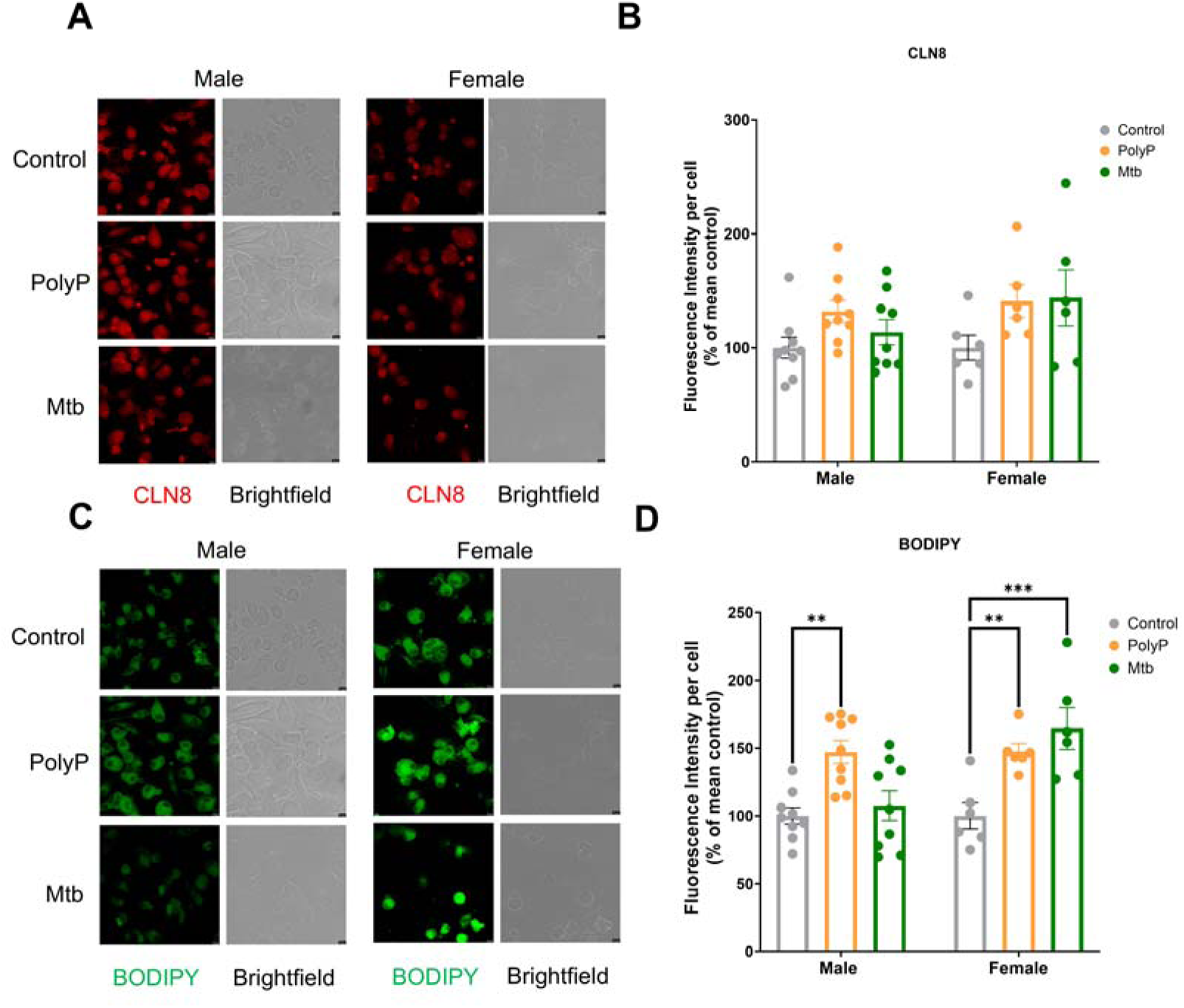
CLN8 immunofluorescence and BODIPY neutral-lipid staining in human macrophages. (**A–D**) Human macrophages from male and female donors were left untreated (Control), treated with 15 µg/mL polyP, or infected with Mtb at an MOI of 5 for 24 hours, fixed, and stained with anti-CLN8 antibodies (**A**) or BODIPY 493/503 (**C**). Fluorescence images for CLN8 (red) (**A**) or BODIPY (green) (**C**) are on the left, and brightfield images are on the right for each treatment condition. Representative images from six independent experiments (three male and three female donors) are shown. Bars are 10 µm. (**B and D**) Mean fluorescence intensity per cell for CLN8 (B) and BODIPY (**D**) was measured from images as in A and C across two regions of interest (ROIs) per female donor and three ROIs per male donor. Individual data points represent ROIs. ROI values were normalized to the average of the untreated-control ROI values from the same donor and expressed as a percentage. Bars show mean ± SEM of donor-level averages from three male and three female donors. **p < 0.01; ***p < 0.001 compared to the control (two-way analysis of variance with Dunnett’s test).

To determine whether polyP or Mtb affected macrophage lipid accumulation, neutral lipid droplets were stained with BODIPY 493/503 and quantified by confocal microscopy (41) (**Figure 6C**). PolyP increased BODIPY fluorescence in macrophages from both male and female donors compared with untreated controls (**Figure 6D**). Mtb also increased BODIPY fluorescence in macrophages from female donors but did not significantly change BODIPY fluorescence in macrophages from male donors (**Figure 6D**). Together, these results indicate that polyP promotes neutral lipid accumulation in macrophages from both sexes, whereas the lipid response to Mtb was detected only in macrophages from female donors.

## Discussion

While broad macrophage transcriptional responses to Mtb infection are well-documented (14, 42), how Mtb-derived polyP specifically might modulate this transcriptional landscape remains unclear. Our findings show that Mtb elicited a substantially larger transcriptional response in female donors than in male donors, with most differentially expressed mRNAs downregulated in the female group. Both Mtb and exogenous polyP selectively altered levels of transcripts associated with phagosome maturation, endolysosomal trafficking, and immune receptor signaling and promoted neutral lipid accumulation. These findings suggest that bacterial polyP may function as a chemical cue that alters host trafficking and metabolic pathways to favor pathogen survival.

We previously observed that uninfected female macrophages have higher basal proteasome activity than male macrophages (13). The higher baseline levels of many Mtb- or polyP-responsive mRNAs in uninfected female macrophages compared to male macrophages provide a wider range for transcriptional downregulation during infection. However, DEG counts from separate analyses of males and females in this small cohort do not, by themselves, show that treatment responses differ by sex.

Compared to Mtb infection, exogenous polyP altered levels of a much smaller set of mRNAs. Because Mtb presents cell-wall lipids, proteins, and secreted effectors in addition to extracellular polyP (43–45), Mtb likely triggers multiple polyP-independent host pathways that contribute to a broader transcriptional response than polyP alone.

Enrichment of gene ontology terms related to cytoplasmic translation, ribosomal subunits, and adaptive immune responses following polyP treatment or Mtb infection provides a potential molecular context for our previous observations that polyP and Mtb inhibit macrophage proteasome activity and proinflammatory polarization (13). However, the present pathway-enrichment results do not show impaired antigen presentation or identify the transcriptional mechanism responsible for reduced proteasome activity. Terms associated with T-cell and NK-cell functions may also reflect residual non-myeloid leukocytes remaining after adherence purification of PBMCs (46).

Although RNA-seq highlighted *PSMB9*, *HLA-DPA1*, *PLEKHM1*, and *SCARF1* as candidate genes, their RNA-seq expression patterns were not reproduced by qRT-PCR. Expression of *HLA-DPA1* in females and *PLEKHM1* and *PSMB9* in males varied substantially among individual donors, which may contribute to the divergence between the DESeq2 results and the donor-normalized qRT-PCR measurements. We therefore interpret these four genes as exploratory candidates. For the validated candidates, the broader detection of *CORO1A*, *S1PR1*, and *INPP4B* downregulation by qRT-PCR may reflect the genome-wide multiple-testing correction and fold-change threshold applied to RNA-seq.

Coronin 1 (*CORO1A*) is a cytoskeleton-associated protein that transiently associates with phagosomes to regulate actin dynamics (28, 47), and Mtb and extracellular polyP cause coronin 1 to be retained on phagosomes, which may contribute to inhibition of phagosome maturation (11, 48). The observed downregulation of *CORO1A* may represent a compensatory negative feedback response to limit further coronin 1 accumulation following its retention on phagosomes.

Sphingosine-1-phosphate (S1P) signaling stimulates actin assembly on phagosomes, promotes phagolysosomal fusion, and enhances the intracellular killing of mycobacteria, whereas Mtb inhibits sphingosine kinase activity to block phagosome maturation (49–52). The observed downregulation of *S1PR1* by polyP and Mtb may alter macrophage responses to S1P.

Inositol polyphosphate-4-phosphatase type II B (*INPP4B*) converts phosphatidylinositol-3,4-bisphosphate into phosphatidylinositol-3-phosphate to promote PI3K-dependent late endosome formation and endolysosomal trafficking in breast cancer cells (31). We previously found that polyP reduces macrophage levels of vacuolar protein-sorting 34 (VPS34) and early endosomal antigen 1 (EEA1), which are necessary for phagosome acidification and maturation (11), and that pharmacological inhibition of IP6K- and mTORC1-associated pathways enhances macrophage bacterial killing (13). Downregulation of *INPP4B* by polyP and Mtb may also impair endolysosomal trafficking by reducing INPP4B-mediated phosphoinositide conversion.

The increase in CLN8 mRNA, which encodes a protein that facilitates the transport of lysosomal enzymes and is required for lysosome biogenesis (29), in Mtb-infected female macrophages may represent a compensatory response to inhibition of lysosome activity (11), although no corresponding increase in CLN8 protein immunofluorescence was detected at 24 hours. *CLN8* translation rates and protein turnover were not measured, leaving the basis of the mRNA–protein discrepancy unresolved.

In addition to inhibiting lysosomal function, Mtb alters host lipid metabolism to induce lipid-laden foamy macrophages that support intracellular Mtb persistence (39, 40). Because lysosomes and autophagy degrade intracellular lipid droplets (53), and polyP inhibits lysosome activity (11), the polyP-induced neutral lipid accumulation in macrophages from both sexes may reflect reduced lysosomal lipid degradation. A significant Mtb-associated increase in neutral lipid accumulation in female but not male macrophages does not, by itself, show a sex-by-treatment difference.

This study has several limitations. First, the small cohort of three male and three female healthy donors per assay group restricts the statistical power and generalizability of the transcriptomic and phenotypic observations. Second, we used an attenuated leucine and pantothenate auxotroph of Mtb (54), and macrophage gene expression differs between infections with virulent and attenuated strains (14). Third, our bulk RNA-seq approach measures population-average transcript levels at a single 24-hour time point and cannot resolve temporal or single-cell variation (55–57). Finally, because polyP signaling can depend on polymer chain length and local concentration (58), our use of a single concentration (15 µg/mL) of long-chain polyP (∼700 phosphate units) does not capture potential chain-length- or dose-dependent transcriptional effects.

Targeted knockdowns or knockouts of the validated candidate genes will help determine whether these genes contribute to polyP-mediated macrophage dysfunction. Building upon our recent findings that pharmacological targeting of host pathways can restore macrophage bactericidal activity (13), testing these candidate pathways may advance host-directed therapeutic strategies against tuberculosis.

## Materials and methods

### Human cell culture

Human peripheral blood was collected from six healthy volunteer blood donors (three males and three females, aged 18 to 35 years) for RNA sequencing and qRT-PCR experiments and six additional healthy blood donors (three males and three females, aged 18 to 35 years) for immunofluorescence and neutral-lipid microscopy. All donors provided written consent, with specific approval from the institutional review board at Texas A&M for blood collected there and the institutional review board at the University of Southern Mississippi for blood collected there. Peripheral blood mononuclear cells (PBMCs) were isolated and purified as previously described (59). A total of 20 × 10^6^ PBMCs were resuspended in 20 mL of complete RPMI culture medium consisting of Roswell Park Memorial Institute (RPMI) 1640 medium (12-167F, Lonza, Walkersville, MD, USA) containing 10% (vol/vol) bovine calf serum (2100-500, VWR Life Science Seradigm, Radnor, PA, USA), 2 mM L-glutamine (Lonza), and 50 ng/mL recombinant human granulocyte-macrophage colony-stimulating factor (GM-CSF; 572903, BioLegend, San Diego, CA, USA) (17). For RNA-sequencing and qRT-PCR experiments, PBMCs were seeded at a density of 1 × 10^6^ cells/mL (5 mL per well) in type 3516 6-well, tissue-culture-treated plates (Corning, Big Flats, NY, USA). For immunofluorescence and lipid-droplet microscopy assays, PBMCs were seeded at a density of 1 × 10^5^ cells/well in 100 µL of complete RPMI culture medium in type 353219 96-well, black/clear, tissue-culture-treated, glass-bottom plates (Corning, Big Flats, NY, USA). Cells were incubated at 37 °C in a humidified incubator with 5% CO_2_ for 6 days to allow differentiation of monocytes into macrophages, with fresh complete RPMI medium replenished on day 6 (11).

### Bacterial cell culture

The attenuated Biosafety Level 2 (BSL-2) Mtb Δ*leuD*Δ*panCD* double auxotroph of *Mycobacterium tuberculo*sis H37Rv (Mtb; a gift from Dr. Jim Sacchettini, Department of Biochemistry and Biophysics, Texas A&M University, College Station, TX, USA) (54) was grown as previously described (60–62) in Middlebrook 7H9 broth (BD, Sparks, MD, USA) or on Middlebrook 7H10 agar (BD) supplemented with 0.5% (vol/vol) glycerol (VWR Life Science Seradigm), 0.05% (vol/vol) Tween 80 (MP Biomedicals, Solon, OH, USA), 10% (vol/vol) Middlebrook Oleic Albumin Dextrose Catalase (OADC) Enrichment (BD), 50 μg/mL L-leucine (VWR Life Science Seradigm), and 50 μg/mL calcium pantothenate (Beantown Chemical, Hudson, NH, USA). Liquid cultures were grown at 37 °C in 50 mL conical tubes on a STR200-V variable angle tube rotator (Southwest Science, Roebling, NJ) in a humidified incubator for 1 to 2 weeks until log phase (OD_600_ of 0.6). The 600 nm optical density of 100 µL of the culture was measured with a Synergy Mx monochromator microplate reader (Biotek, Winooski, VT). Once an OD_600_ of 0.6 was reached, the cells were washed with complete RPMI medium supplemented with L-leucine, pantothenate and GM-CSF by centrifugation at 12,000 x g for 2 minutes in a microcentrifuge tube, and resuspended in 1 mL of RPMI complete medium and diluted to OD_600_ of 0.02 to be used in the assays.

### PolyP preparation

Long-chain sodium polyphosphate (polyP; p700, average chain length ∼700 phosphate units; Cat# EUI002, Kerafast, Inc., Boston, MA, USA) was reconstituted in sterile purified water to a stock concentration of 102 mg/mL polyP according to the manufacturer’s instructions and diluted in culture medium to a final working concentration of 15 µg/mL (11).

### Macrophage infection, polyP treatment, and RNA extraction

On day 7 of macrophage differentiation, the culture supernatant and loosely adhered cells were removed, and 5 mL of complete RPMI medium supplemented with L-leucine, pantothenate and GM-CSF, as described above, were added to each well of the 6-well plates. For polyP-treated wells, polyP was added and plates were incubated for 30 minutes at 37 °C with 5 % CO_2_. For Mtb-infected wells, log-phase Mtb bacteria were added at an OD_600_ of 0.02, corresponding to a multiplicity of infection (MOI) of ∼5 bacteria per macrophage, and mixed gently. Control wells were left untreated and uninfected. The plates were incubated at 37 °C in 5% CO_2_ for 2 hours to allow phagocytosis. After 2 hours, the supernatant medium was removed, and each well was washed with 5 mL of pre-warmed (37 °C) phosphate-buffered saline (PBS; Gibco, Thermo Fisher Scientific, Waltham, MA, USA) to remove un-ingested extracellular bacteria. Fresh medium supplemented with 200 µg/mL gentamicin (Sigma-Aldrich, St. Louis, MO, USA) was added to kill remaining extracellular bacteria, and the plates were incubated for an additional 2 hours at 37 °C. Cells were then washed twice with pre-warmed PBS, replenished with fresh medium (with polyP maintained in the polyP-treated condition), and incubated for 24 hours at 37 °C in 5 % CO_2_. At 24 hours, the culture supernatant was discarded, and total cellular RNA was isolated in a 100 µL elution volume using a Quick-RNA Miniprep Kit (Cat# R1054, Zymo Research, Irvine, CA, USA) according to the manufacturer’s instructions. RNA concentrations and quality were assessed following the manufacturer’s instructions, and 20 ng of total RNA per sample was submitted to the Texas A&M Institute for Genome Sciences and Society (TIGSS) Molecular Genomics Core for low-input cDNA library preparation using the SMART-Seq mRNA LP (with UMIs) kit (Cat# 634762, Takara Bio USA, Inc., San Jose, CA, USA) and Unique Dual Index Kit (1–24) (Cat# 634756, Takara Bio USA, Inc.) followed by next-generation RNA sequencing on an Illumina NextSeq 2000 platform.

### RNA sequencing, differential gene expression, and pathway enrichment analyses

Raw sequencing read libraries for each sample (n = 18; 3 male and 3 female donors across control, polyP, and Mtb conditions) were quality-checked using FastQC (63) and trimmed to remove low-quality base calls (Phred score < 20, -q 20), poly(A) tails (-a), and residual adapter or barcode sequences using Trim Galore, a wrapper around Cutadapt (64). Trimmed RNA-seq reads were mapped and quantified against the human reference transcriptome (GRCh38) using Salmon (version 5) (65). All downstream bioinformatic and statistical analyses were performed in the R programming environment. Transcript-level abundance estimates were imported and summarized to a gene-level count matrix using the tximport R package (66). Principal component analysis (PCA) (18) and hierarchical clustering of sample-to-sample Euclidean distances were performed on variance-stabilized transformed counts to evaluate global sample clustering by donor sex, individual identity, and treatment using pheatmap R package (67). Differential gene expression analysis was conducted using the DESeq2 R package (68) with the design formula ∼ Batch + Sex + Phenotype + Sex: Phenotype to model batch, donor sex (male and female), and treatment phenotype (control, polyP, and Mtb), as well as sex-by-phenotype interactions. For the combined sex-averaged analysis, a numeric contrast vector weighing male-specific and female-specific effects equally was applied. Differentially expressed genes (DEGs) were defined using the DESeq2 Wald test as genes exhibiting an absolute log2 fold change > 1 and a Benjamini–Hochberg-adjusted *P* value (*P*_adj_ < 0.05). Volcano plots and row-scaled (*Z*-score) hierarchical clustering heatmaps were generated using the EnhancedVolcano (69) and ComplexHeatmap (70) R packages, respectively. Gene Ontology (GO) enrichment analysis (71) for biological processes (BP), cellular components (CC), and molecular functions (MF) was performed on all genes significantly altered by each treatment (*P*_adj_ < 0.05) using the topGO R package (72) and human genome annotation package org.Hs.eg.db. Enriched GO pathways were defined as those with *P*_adj_ < 0.05.

### Validation of candidate genes by qRT-PCR

Expression of the selected genes were validated by quantitative reverse transcription-PCR (qRT-PCR). Complementary DNA (cDNA) was synthesized from 150 ng of total RNA per sample using the Maxima H Minus First Strand cDNA Synthesis Kit with dsDNase (Cat# K1682, Thermo Fisher Scientific, Waltham, MA, USA) according to the manufacturer’s protocol. Quantitative PCR reactions were performed using Fast SYBR Green Master Mix (Applied Biosystems, Thermo Fisher Scientific) on a CFX Opus 96 Real-Time PCR System (Bio-Rad Laboratories, Hercules, CA, USA). Amplification was carried out for 30 cycles followed by melting-curve analysis to verify primer specificity. Gene-specific primers were obtained from previously published studies (34, 73–76) or designed using the NCBI Primer-BLAST tool **(Table S1)**. Relative transcript expression was calculated using the comparative *C_T_* (2 ΔΔ*C*_T_) method (77) normalized to the reference gene glyceraldehyde-3-phosphate dehydrogenase (*GAPDH*) (78). For each donor and sex, relative expression values in polyP-treated and Mtb-infected macrophages were expressed as a percentage of the corresponding untreated control, which was set to 100%.

### Immunofluorescence and neutral-lipid droplet microscopy

Macrophages differentiated in 96-well glass-bottom plates (1 × 10 cells per well in 100 μL complete RPMI medium) were left untreated (control), treated with polyP, or infected with Mtb for 24 hours as described above. At 24 hours, cells were gently washed with PBS and fixed with 4% (wt/vol) paraformaldehyde (PFA; Thermo Fisher Scientific) in PBS for 10 minutes at room temperature (79). Fixed cells were washed with PBS containing 0.1% (vol/vol) Tween 20 (Fisher Scientific, Pittsburgh, PA, USA) (PBST), permeabilized with 0.1% (vol/vol) Triton X-100 (Thermo Fisher Scientific) in PBS for 5 minutes at room temperature, and washed twice with PBS. Nonspecific binding sites were blocked with 0.1% (wt/vol) bovine serum albumin (BSA; Fisher BioReagents, Fair Lawn, NJ, USA) in PBS for 1 hour at 4°C. Cells were washed once with PBS and incubated overnight at 4°C in the dark with rabbit polyclonal anti-CLN8 primary antibody (Cat# PA5-113894, Invitrogen, Thermo Fisher Scientific) diluted to 0.5 µg/mL in PBST. Following three 5-minute washes with PBST, cells were incubated for 1 hour at room temperature in the dark with PBST containing 1 µg/mL Alexa Fluor 568-conjugated goat anti-rabbit IgG secondary antibody (Cat# A-11011, Invitrogen, Thermo Fisher Scientific), 10 µM BODIPY 493/503 (4,4-difluoro-1,3,5,7,8-pentamethyl-4-bora-3a,4a-diaza-s-indacene; Cat# D3922, Invitrogen, Thermo Fisher Scientific) to stain neutral-lipid droplets (41), and 10 µg/mL DAPI (4’,6-diamidino-2-phenylindole; Cat# D1306, Invitrogen, Thermo Fisher Scientific) to stain nuclei. Cells were washed three times with PBST (5 minutes per wash) and submerged in 200 μL of PBS per well for imaging.

### Confocal microscopy and image quantification

Stained macrophages were imaged using a Leica confocal laser-scanning microscope (Leica Microsystems, Wetzlar, Germany) equipped with a 63X oil-immersion objective. For each donor and treatment condition, three images were acquired from randomly selected fields of view using identical laser power, detector gain, and exposure settings across all samples. Fluorescence intensities for CLN8 and BODIPY 493/503 were quantified using Fiji (ImageJ; National Institutes of Health, Bethesda, MD, USA) software (80). Images were converted to 8-bit format with a uniform threshold set to 45, and background-subtracted integrated fluorescence intensity density was measured and divided by the total number of cells per field to obtain the mean fluorescence intensity per cell. For each donor and sex, the mean fluorescence intensity per cell was measured in two regions of interest (ROIs) for female donors and three ROIs for male donors. Each ROI value was normalized to the average of the untreated-control ROI values from the same donor and expressed as a percentage, with the control average set to 100%.

### Statistical analysis

Statistical analyses for RNA-seq differential expression and GO enrichment were performed in R using the DESeq2 and topGO packages as described above, and statistical analyses for qRT-PCR and confocal microscopy assays were performed using Prism 11.0.2 (GraphPad Software, San Diego, CA, USA) and the tests indicated in the figure legends. *P* < 0.05 was considered significant.

## Data availability

The RNA-sequencing data generated in this study have been deposited in the NCBI Gene Expression Omnibus (GEO) under accession number (GSE355811). All other study data are included in the article and supplemental material. Further information and requests for resources and reagents may be directed to R.R.

## Acknowledgments

The authors are thankful to the volunteers who donated blood and the phlebotomy staff at the Texas A&M Beutel Student Health Center and the University of Southern Mississippi Medical Lab Science. Use of the Texas A&M Institute for Genome Science & Society (TIGSS) Molecular Genomics Core and Texas A&M High-Performance Research Computing is acknowledged, and we thank Devon J. Boland for RNA-seq data analysis assistance. We thank Takara Bio USA, Inc. for providing the SMART-Seq mRNA LP (with UMIs) kit and library preparation support through the TIGSS–Takara Bio Low-Input Sequencing Contest Award. We also thank the Mississippi INBRE Imaging Core Facility at The University of Southern Mississippi for confocal microscopy support.

This work was supported by the National Institutes of Health grant R35GM118355 (to R.H.G.), R16AI192297 (to R.R), an institutional start-up fund from The University of Southern Mississippi (to R.R.), and the Mississippi INBRE Imaging Core Facility at The University of Southern Mississippi, funded by an Institutional Development Award (IDeA) from the National Institute of General Medical Sciences of the National Institutes of Health under grant number P20GM103476.

A.T., M.M., and R.R. designed and performed experiments and analyzed data; A.T. and R.R. drafted the paper; and R.R. and R.H.G. coordinated the study, revised the paper, and acquired funding.

## Conflict of Interest

The authors declare no competing interests.

## AI Usage Policy

Generative AI technologies were used solely to improve the readability and language of the manuscript, and all AI-assisted writing was carefully reviewed and edited by the authors, who take full responsibility for the content of the publication.

## Supplementary figures

**Figure S1.**
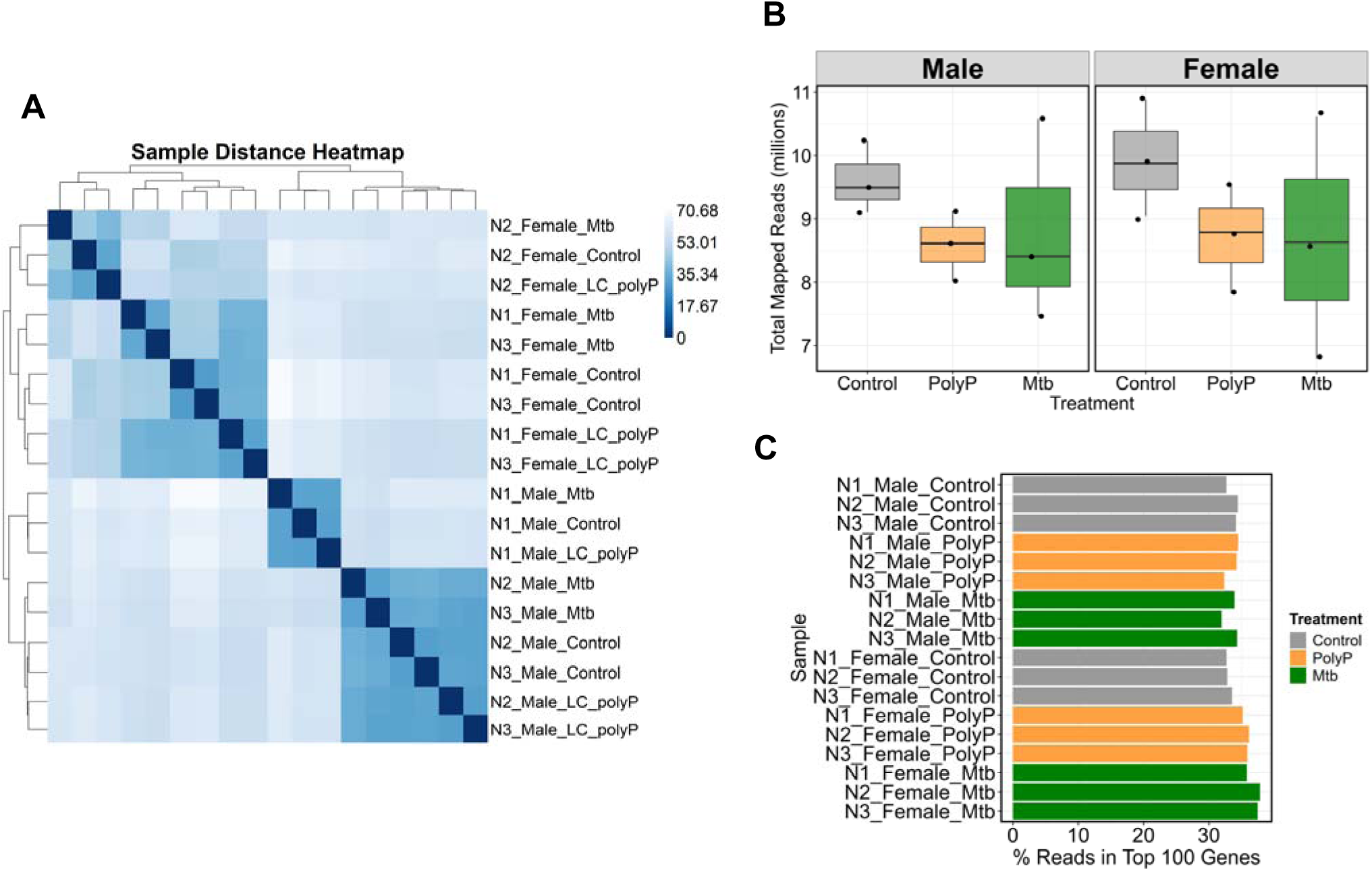
Sample-to-sample distance clustering, sequencing depth, and library complexity. (**A**) Heatmap and hierarchical clustering dendrogram of sample-to-sample Euclidean distances across the 18 macrophage RNA-seq libraries from male and female donors under control, polyP, and Mtb conditions. (**B**) Box plots of total mapped reads (in millions) per sample grouped by treatment condition (Control, gray; PolyP, orange; Mtb, green) in male and female donors. (**C**) Percentage of total mapped reads accounted for by the 100 most highly expressed genes in each sample across control (gray), polyP (orange), and Mtb (green) conditions.

**Figure S2.**
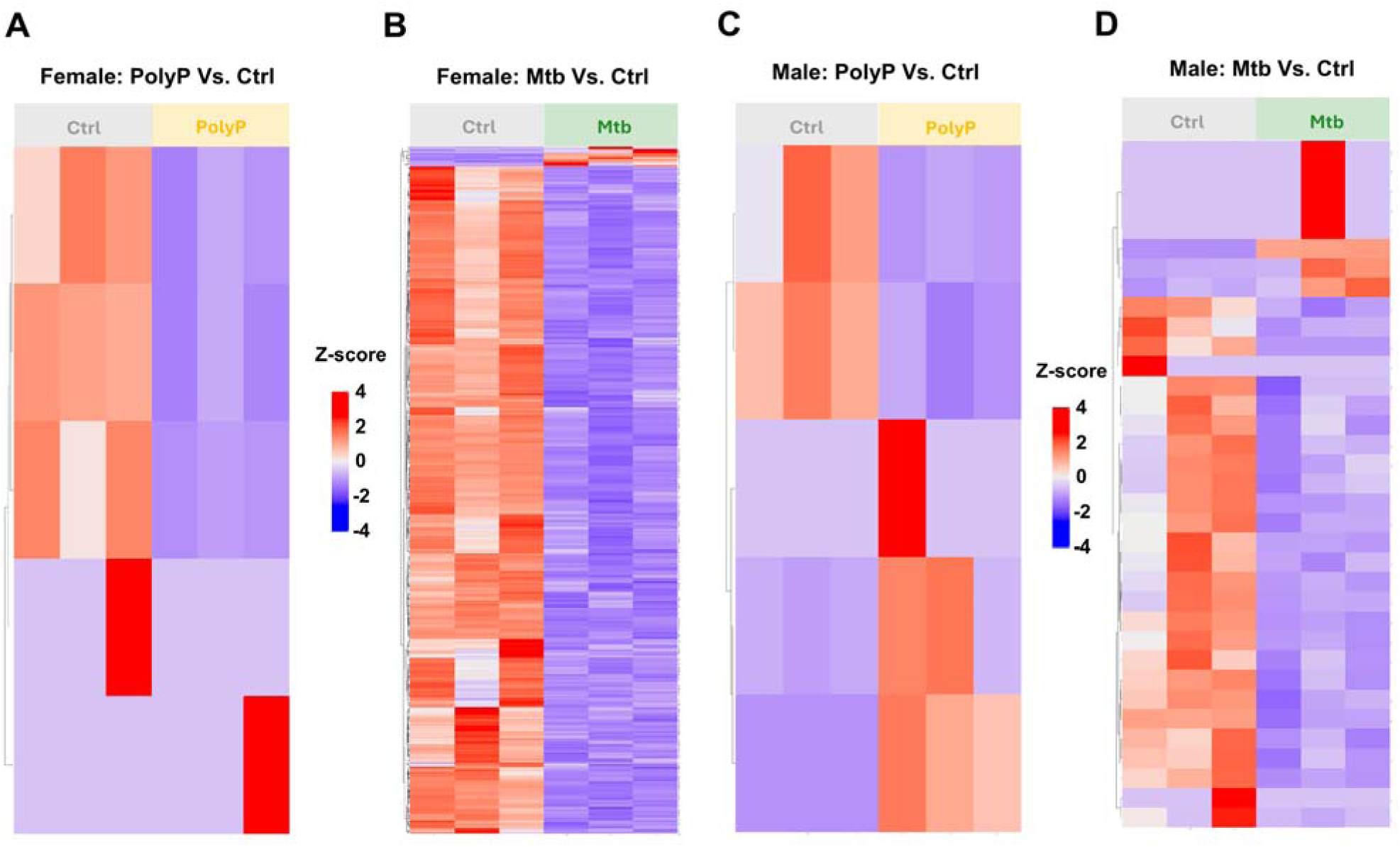
Clustered expression heatmaps in the sex-stratified analysis. (**A and B**) Heatmaps of row-scaled (*Z*-score) expression values for differentially expressed genes comparing polyP-treated vs. control (Ctrl) (**A**) and Mtb-infected vs. control (**B**) macrophages from female donors. (**C and D**) Heatmaps of row-scaled (*Z*-score) expression values for differentially expressed genes comparing polyP-treated vs. control (**C**) and Mtb-infected vs. control (**D**) macrophages from male donors.

**Figure S3.**
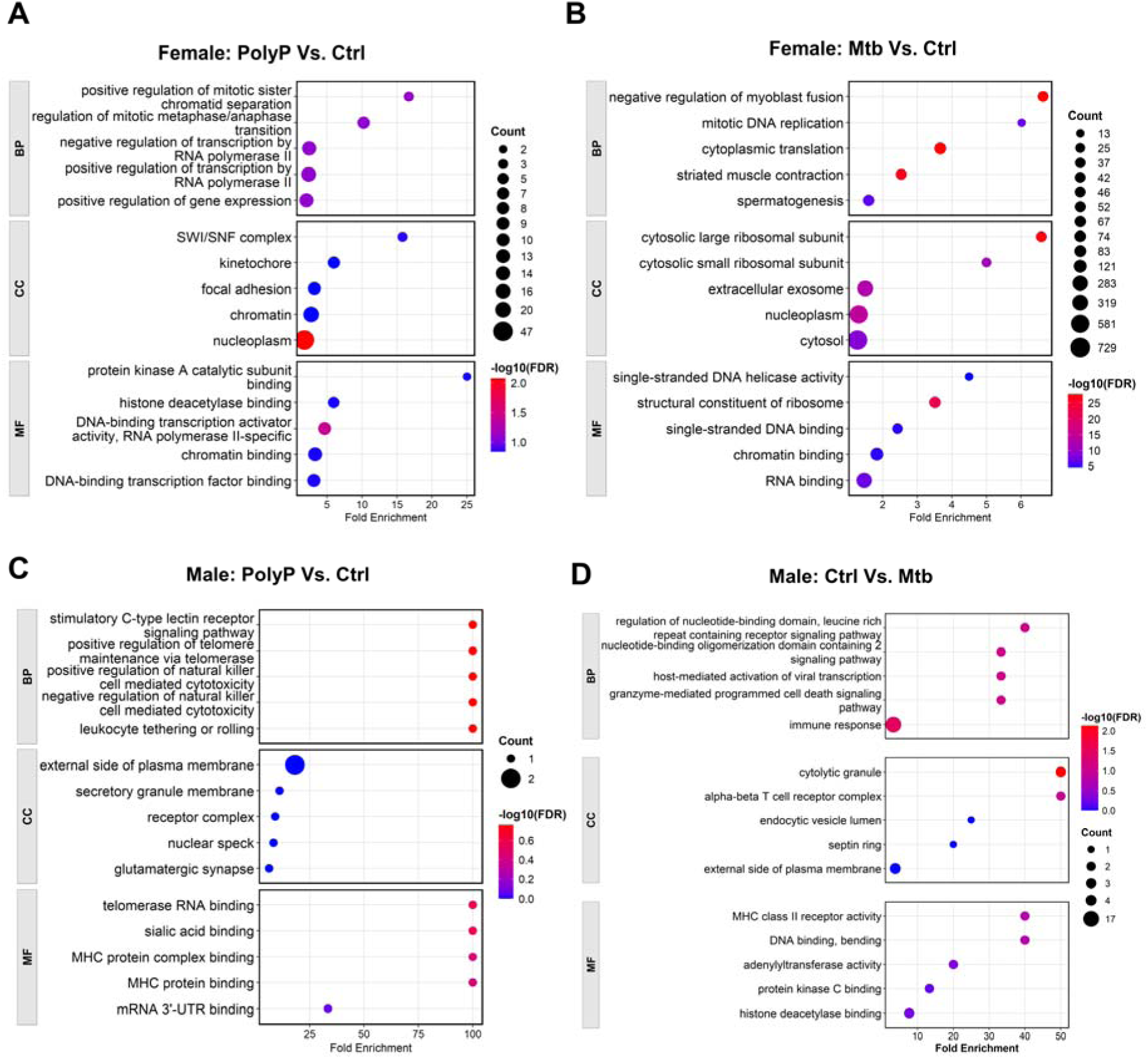
Gene Ontology enrichment in the sex-stratified analysis. (**A and B**) Gene Ontology (GO) enrichment dot plots for biological processes (BP), cellular components (CC), and molecular functions (MF) of genes significantly altered by polyP (**A**) or Mtb (**B**) compared with controls in female donors. (**C and D**) GO enrichment dot plots for biological processes (BP), cellular components (CC), and molecular functions (MF) of genes significantly altered by polyP (**C**) or Mtb (**D**) compared with controls in male donors. In **A, B, C, and D**, the top five enriched terms in each category are plotted by fold enrichment; dot size indicates the number of enriched genes (Count), and dot color indicates –log10(FDR).

**Figure S4.**
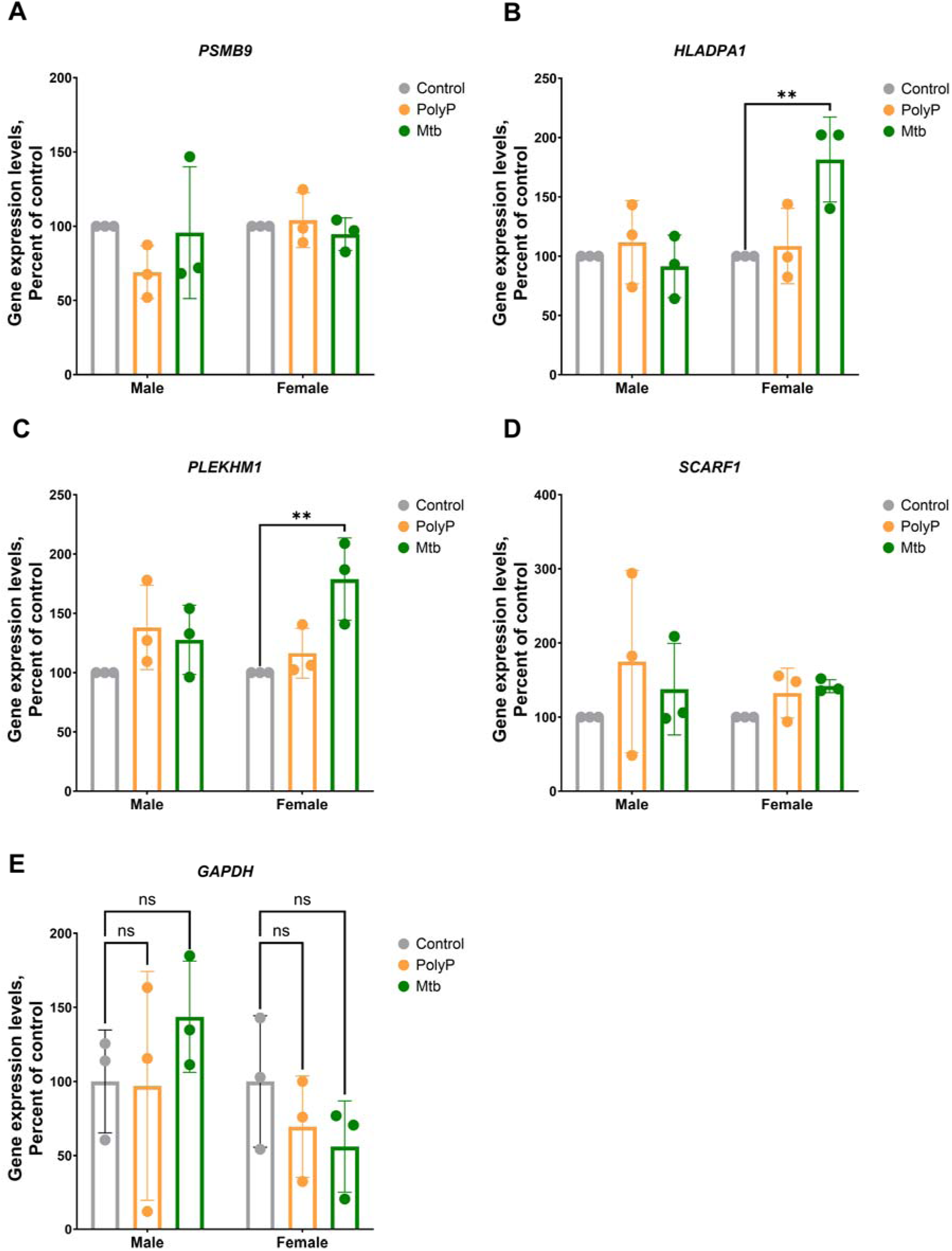
qRT-PCR analysis of additional candidate genes and GAPDH reference expression. (**A–E**) Relative mRNA expression levels of *PSMB9* (**A**), *HLA-DPA1* (**B**), *PLEKHM1* (**C**), *SCARF1* (**D**), and the reference gene *GAPDH* (**E**) in untreated (Control, gray), polyP-treated (PolyP, orange), and Mtb-infected (Mtb, green) macrophages from male and female donors at 24 hours, determined by qRT-PCR. Expression levels in **A–D** were normalized to *GAPDH*. For each experiment with each donor, expression in the control was set to 100%. All values are mean ± SEM of three male and three female donors. *\*\*p < 0.01* compared to the control; ns, not significant (two-way analysis of variance with Dunnett’s test).

**Table S1.**
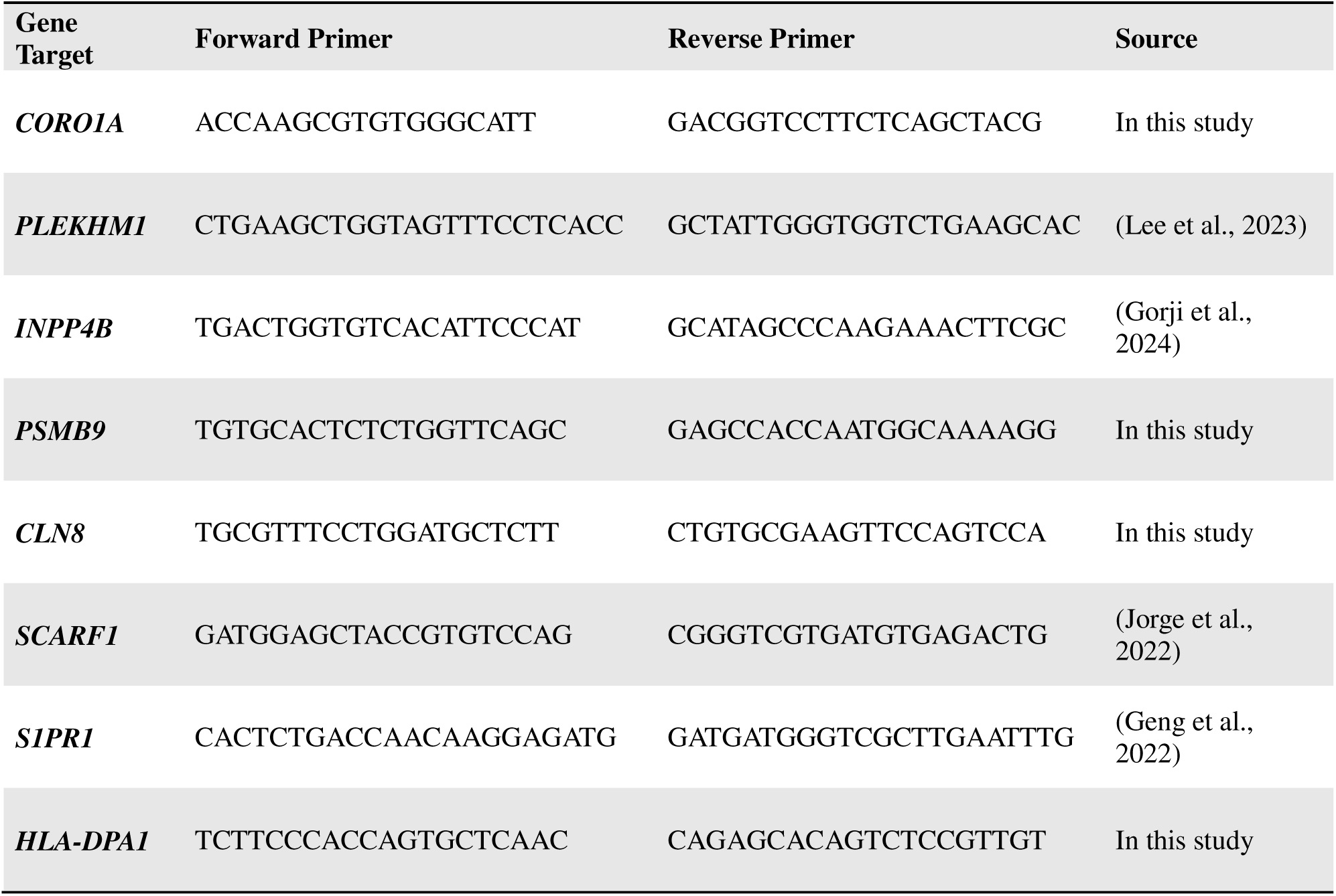
Gene-specific primers obtained from previously published studies or designed using the NCBI Primer-BLAST tool.

## Notes

https://www.ncbi.nlm.nih.gov/geo/query/acc.cgi?acc=GSE355811

### Competing Interest Statement

The authors have declared no competing interest.

## References

1. Y. M. Chuang, D. A. Belchis, P. C. Karakousis, The polyphosphate kinase gene ppk2 is required for Mycobacterium tuberculosis inorganic polyphosphate regulation and virulence. mBio 4, e00039–00013 (2013).

2. K. S. Kim, N. N. Rao, C. D. Fraley, A. Kornberg, Inorganic polyphosphate is essential for long-term survival and virulence factors in Shigella and Salmonella spp. Proc Natl Acad Sci U S A 99, 7675–7680 (2002).

3. N. N. Rao, M. R. Gomez-Garcia, A. Kornberg, Inorganic polyphosphate: essential for growth and survival. Annu Rev Biochem 78, 605–647 (2009).

4. M. H. Rashid et al., Polyphosphate kinase is essential for biofilm development, quorum sensing, and virulence of Pseudomonas aeruginosa. Proc Natl Acad Sci U S A 97, 9636–9641 (2000).

5. R. S. Flannagan, G. Cosio, S. Grinstein, Antimicrobial mechanisms of phagocytes and bacterial evasion strategies. Nat Rev Microbiol 7, 355–366 (2009).

6. I. Jutras, M. Desjardins, Phagocytosis: at the crossroads of innate and adaptive immunity. Annu Rev Cell Dev Biol 21, 511–527 (2005).

7. C. Nathan, M. U. Shiloh, Reactive oxygen and nitrogen intermediates in the relationship between mammalian hosts and microbial pathogens. Proc Natl Acad Sci U S A 97, 8841–8848 (2000).

8. WHO (2025) Global tuberculosis report: . (Geneva: World Health Organization; 2025. Licence: CC BY-NC-SA 3.0 IGO).

9. C. J. Queval, R. Brosch, R. Simeone, The Macrophage: A Disputed Fortress in the Battle against Mycobacterium tuberculosis. Front Microbiol 8, 2284 (2017).

10. D. M. Murphy et al., Human Macrophages Activate Bystander Neutrophils’ Metabolism and Effector Functions When Challenged with Mycobacterium tuberculosis. Int J Mol Sci 25 (2024).

11. R. Rijal, L. A. Cadena, M. R. Smith, J. F. Carr, R. H. Gomer, Polyphosphate is an extracellular signal that can facilitate bacterial survival in eukaryotic cells. Proceedings of the National Academy of Sciences 117, 31923–31934 (2020).

12. R. J. Rahman et al., Polyphosphate uses mTOR, pyrophosphate, and Rho GTPase components to potentiate bacterial survival in Dictyostelium. mBio 14, e0193923 (2023).

13. R. Rijal, R. H. Gomer, Pharmacological inhibition of host pathways enhances macrophage killing of intracellular bacterial pathogens. Microbiol Spectr 14, e0216325 (2026).

14. J. Lee et al., Characterisation of genes differentially expressed in macrophages by virulent and attenuated Mycobacterium tuberculosis through RNA-Seq analysis. Sci Rep 9, 4027 (2019).

15. J. Dibbern, L. Eggers, B. E. Schneider, Sex differences in the C57BL/6 model of Mycobacterium tuberculosis infection. Sci Rep 7, 10957 (2017).

16. H. M. Rickman et al., Sex differences in the risk of Mycobacterium tuberculosis infection: a systematic review and meta-analysis of population-based immunoreactivity surveys. Lancet Public Health 10, e588–e598 (2025).

17. I. Ushach, A. Zlotnik, Biological role of granulocyte macrophage colony-stimulating factor (GM-CSF) and macrophage colony-stimulating factor (M-CSF) on cells of the myeloid lineage. J Leukoc Biol 100, 481–489 (2016).

18. M. Ringner, What is principal component analysis? Nat Biotechnol 26, 303–304 (2008).

19. D. A. Patten, SCARF1: a multifaceted, yet largely understudied, scavenger receptor. Inflamm Res 67, 627–632 (2018).

20. T. Onodera et al., Btbd7 regulates epithelial cell dynamics and branching morphogenesis. Science 329, 562–565 (2010).

21. M. A. Vostakolaei et al., Hsp70 in cancer: A double agent in the battle between survival and death. J Cell Physiol 236, 3420–3444 (2021).

22. A. Ivetic, H. L. Hoskins Green, S. J. Hart, L-selectin: A Major Regulator of Leukocyte Adhesion, Migration and Signaling. Front Immunol 10, 1068 (2019).

23. J. G. Cyster, S. R. Schwab, Sphingosine-1-phosphate and lymphocyte egress from lymphoid organs. Annu Rev Immunol 30, 69–94 (2012).

24. E. Bongen, F. Vallania, P. J. Utz, P. Khatri, KLRD1-expressing natural killer cells predict influenza susceptibility. Genome Med 10, 45 (2018).

25. H. Liang, X. Liang, M. Zheng, S. Wang, Elucidating the role of KLRD1 in coronary atherosclerosis: harnessing bioinformatics and machine learning to advance understanding. J Cardiothorac Surg 20, 248 (2025).

26. L. Santiago, G. Daniels, D. Wang, F. M. Deng, P. Lee, Wnt signaling pathway protein LEF1 in cancer, as a biomarker for prognosis and a target for treatment. Am J Cancer Res 7, 1389–1406 (2017).

27. M. B. Eisen, P. T. Spellman, P. O. Brown, D. Botstein, Cluster analysis and display of genome-wide expression patterns. Proc Natl Acad Sci U S A 95, 14863–14868 (1998).

28. M. Yan, R. F. Collins, S. Grinstein, W. S. Trimble, Coronin-1 function is required for phagosome formation. Mol Biol Cell 16, 3077–3087 (2005).

29. A. di Ronza et al., CLN8 is an endoplasmic reticulum cargo receptor that regulates lysosome biogenesis. Nat Cell Biol 20, 1370–1377 (2018).

30. R. Marwaha et al., The Rab7 effector PLEKHM1 binds Arl8b to promote cargo traffic to lysosomes. J Cell Biol 216, 1051–1070 (2017).

31. S. J. Rodgers et al., INPP4B promotes PI3Kalpha-dependent late endosome formation and Wnt/beta-catenin signaling in breast cancer. Nat Commun 12, 3140 (2021).

32. M. Kasahara, M. F. Flajnik, Origin and evolution of the specialized forms of proteasomes involved in antigen presentation. Immunogenetics 71, 251–261 (2019).

33. Y. Yamashita et al., HLA-DP(84Gly) constitutively presents endogenous peptides generated by the class I antigen processing pathway. Nat Commun 8, 15244 (2017).

34. Y. Wei et al., Integrating single-cell RNA-seq and bulk RNA-seq to identify hub pathways and diagnostic biomarkers in active pulmonary tuberculosis. BMC Infect Dis 26 (2026).

35. S. K. Garg et al., Sphingosine 1-phosphate induces antimicrobial activity both in vitro and in vivo. J Infect Dis 189, 2129–2138 (2004).

36. S. J. Rodgers et al., INPP4B promotes PI3Kalpha-dependent late endosome formation and Wnt/beta-catenin signaling in breast cancer. Nat Commun 12, 3140 (2021).

37. S. Seto, K. Tsujimura, Y. Koide, Coronin-1a inhibits autophagosome formation around Mycobacterium tuberculosis-containing phagosomes and assists mycobacterial survival in macrophages. Cell Microbiol 14, 710–727 (2012).

38. L. Lonka, A. Kyttala, S. Ranta, A. Jalanko, A. E. Lehesjoki, The neuronal ceroid lipofuscinosis CLN8 membrane protein is a resident of the endoplasmic reticulum. Hum Mol Genet 9, 1691–1697 (2000).

39. D. G. Russell, P. J. Cardona, M. J. Kim, S. Allain, F. Altare, Foamy macrophages and the progression of the human tuberculosis granuloma. Nat Immunol 10, 943–948 (2009).

40. P. Peyron et al., Foamy macrophages from tuberculous patients’ granulomas constitute a nutrient-rich reservoir for M. tuberculosis persistence. PLoS Pathog 4, e1000204 (2008).

41. T. Govender, L. Ramanna, I. Rawat, F. Bux, BODIPY staining, an alternative to the Nile Red fluorescence method for the evaluation of intracellular lipids in microalgae. Bioresour Technol 114, 507–511 (2012).

42. W. Pu et al., Comparative transcriptomic analysis of THP-1-derived macrophages infected with Mycobacterium tuberculosis H37Rv, H37Ra and BCG. J Cell Mol Med 25, 10504–10520 (2021).

43. S. L. Sampson, Mycobacterial PE/PPE proteins at the host-pathogen interface. Clin Dev Immunol 2011, 497203 (2011).

44. S. Upadhyay, E. Mittal, J. A. Philips, Tuberculosis and the art of macrophage manipulation. Pathog Dis 76 (2018).

45. C. Ghazaei, Mycobacterium tuberculosis and lipids: Insights into molecular mechanisms from persistence to virulence. J Res Med Sci 23, 63 (2018).

46. M. C. Nielsen, M. N. Andersen, H. J. Moller, Monocyte isolation techniques significantly impact the phenotype of both isolated monocytes and derived macrophages in vitro. Immunology 159, 63–74 (2020).

47. S. Itoh et al., The role of protein kinase C in the transient association of p57, a coronin family actin-binding protein, with phagosomes. Biol Pharm Bull 25, 837–844 (2002).

48. G. Ferrari, H. Langen, M. Naito, J. Pieters, A Coat Protein on Phagosomes Involved in the Intracellular Survival of Mycobacteria. Cell 97, 435–447 (1999).

49. E. Anes et al., Selected lipids activate phagosome actin assembly and maturation resulting in killing of pathogenic mycobacteria. Nat Cell Biol 5, 793–802 (2003).

50. Z. A. Malik et al., Cutting edge: Mycobacterium tuberculosis blocks Ca2+ signaling and phagosome maturation in human macrophages via specific inhibition of sphingosine kinase. J Immunol 170, 2811–2815 (2003).

51. I. Treede et al., Anti-inflammatory effects of phosphatidylcholine. J Biol Chem 282, 27155–27164 (2007).

52. A. Weigert, N. Weis, B. Brune, Regulation of macrophage function by sphingosine-1-phosphate. Immunobiology 214, 748–760 (2009).

53. R. Singh et al., Autophagy regulates lipid metabolism. Nature 458, 1131–1135 (2009).

54. S. L. Sampson et al., Protection elicited by a double leucine and pantothenate auxotroph of Mycobacterium tuberculosis in guinea pigs. Infect Immun 72, 3031–3037 (2004).

55. A. K. Shalek et al., Single-cell transcriptomics reveals bimodality in expression and splicing in immune cells. Nature 498, 236–240 (2013).

56. R. Avraham et al., Pathogen Cell-to-Cell Variability Drives Heterogeneity in Host Immune Responses. Cell 162, 1309–1321 (2015).

57. A. K. Shalek et al., Single-cell RNA-seq reveals dynamic paracrine control of cellular variation. Nature 510, 363–369 (2014).

58. J. Roewe et al., Bacterial polyphosphates interfere with the innate host defense to infection. Nat Commun 11, 4035 (2020).

59. D. Pilling, V. Vakil, R. H. Gomer, Improved serum-free culture conditions for the differentiation of human and murine fibrocytes. J Immunol Methods 351, 62–70 (2009).

60. M. H. Larsen, K. Biermann, S. Tandberg, T. Hsu, W. R. Jacobs, Jr., Genetic Manipulation of Mycobacterium tuberculosis. Curr Protoc Microbiol Chapter 10, Unit 10A 12 (2007).

61. L. S. Qi et al., Repurposing CRISPR as an RNA-guided platform for sequence-specific control of gene expression. Cell 152, 1173–1183 (2013).

62. J. M. Rock et al., Programmable transcriptional repression in mycobacteria using an orthogonal CRISPR interference platform. Nat Microbiol 2, 16274 (2017).

63. S. Andrews (2010) FASTQC. A quality control tool for high throughput sequence data.

64. M. Martin, Cutadapt removes adapter sequences from high-throughput sequencing reads. EMBnet.journal 17 (2011).

65. R. Patro, G. Duggal, M. I. Love, R. A. Irizarry, C. Kingsford, Salmon provides fast and bias-aware quantification of transcript expression. Nat Methods 14, 417–419 (2017).

66. C. Soneson, M. I. Love, M. D. Robinson, Differential analyses for RNA-seq: transcript-level estimates improve gene-level inferences. F1000Res 4, 1521 (2015).

67. R. Kolde, Pheatmap: pretty heatmaps. R package version 1, 726 (2019).

68. M. I. Love, W. Huber, S. Anders, Moderated estimation of fold change and dispersion for RNA-seq data with DESeq2. Genome Biol 15, 550 (2014).

69. K. Blighe, S Rana, and M Lewis (2018) EnhancedVolcano: Publication-ready volcano plots with enhanced colouring and labeling.

70. Z. Gu, R. Eils, M. Schlesner, Complex heatmaps reveal patterns and correlations in multidimensional genomic data. Bioinformatics 32, 2847–2849 (2016).

71. M. Ashburner et al., Gene ontology: tool for the unification of biology. The Gene Ontology Consortium. Nat Genet 25, 25–29 (2000).

72. R. J. Alexa A, topGO: Enrichment Analysis for Gene Ontology. doi:10.18129/B9.bioc.topGO (2026).

73. X. Geng et al., Downregulation of miR-33a/b and miR-181a contributes to recurrent pregnancy loss by upregulating S1PR1 and repressing regulatory T cell differentiation. Placenta 121, 137–144 (2022).

74. M. Gorji, M. A. Farsani, M. Kargar, J. Garavand, M. H. Mohammadi, Investigating the multifaceted cooperation of autophagy, PI3K/AKT signaling pathways, and INPP4B gene in de novo acute myeloid leukemia patients. Curr Res Transl Med 72, 103429 (2024).

75. A. M. Jorge et al., SCARF1-Induced Efferocytosis Plays an Immunomodulatory Role in Humans, and Autoantibodies Targeting SCARF1 Are Produced in Patients with Systemic Lupus Erythematosus. J Immunol 208, 955–967 (2022).

76. J. W. Lee et al., Centrosome clustering control in osteoclasts through CCR5-mediated signaling. Sci Rep 13, 20813 (2023).

77. K. J. Livak, T. D. Schmittgen, Analysis of relative gene expression data using real-time quantitative PCR and the 2(-Delta Delta C(T)) Method. Methods 25, 402–408 (2001).

78. R. D. Barber, D. W. Harmer, R. A. Coleman, B. J. Clark, GAPDH as a housekeeping gene: analysis of GAPDH mRNA expression in a panel of 72 human tissues. Physiol Genomics 21, 389–395 (2005).

79. U. Schnell, F. Dijk, K. A. Sjollema, B. N. Giepmans, Immunolabeling artifacts and the need for live-cell imaging. Nat Methods 9, 152–158 (2012).

80. J. Schindelin et al., Fiji: an open-source platform for biological-image analysis. Nat Methods 9, 676–682 (2012).

